# Predicting RNA Structure and Dynamics with Deep Learning and Solution Scattering

**DOI:** 10.1101/2024.06.08.598075

**Authors:** Edan Patt, Scott Classen, Michal Hammel, Dina Schneidman-Duhovny

**Affiliations:** School of Computer Science and Engineering, The Hebrew University of Jerusalem; Molecular Biophysics and Integrated Bioimaging, Lawrence Berkeley National Laboratory, Berkeley, CA, USA

## Abstract

Advanced deep learning and statistical methods can predict structural models for RNA molecules. However, RNAs are flexible, and it remains difficult to describe their macromolecular conformations in solutions where varying conditions can induce conformational changes. Small-angle X-ray scattering (SAXS) in solution is an efficient technique to validate structural predictions by comparing the experimental SAXS profile with those calculated from predicted structures. There are two main challenges in comparing SAXS profiles to RNA structures: the absence of cations essential for stability and charge neutralization in predicted structures and the inadequacy of a single structure to represent RNA’s conformational plasticity. We introduce Solution Conformation Predictor for RNA (SCOPER) to address these challenges. This pipeline integrates kinematics-based conformational sampling with the innovative deep-learning model, IonNet, designed for predicting Mg^2+^ ion binding sites. Validated through benchmarking against fourteen experimental datasets, SCOPER significantly improved the quality of SAXS profile fits by including Mg^2+^ ions and sampling of conformational plasticity. We observe that an increased content of monovalent and bivalent ions leads to decreased RNA plasticity. Therefore, carefully adjusting the plasticity and ion density is crucial to avoid overfitting experimental SAXS data. SCOPER is an efficient tool for accurately validating the solution state of RNAs given an initial, sufficiently accurate structure and provides the corrected atomistic model, including ions.

The method is available from: https://github.com/dina-lab3d/IonNet

Our pipeline is available for use as a web server: https://bilbomd.bl1231.als.lbl.gov/

**Statement of Significance:** Understanding the behavior of RNA in solution is critical for deciphering its biological functions, yet predicting its macromolecular conformation remains challenging. While advanced computational methods can predict RNA structures, their accuracy in solution is often limited by the absence of stabilizing ions and the failure to account for RNA’s conformational flexibility. This study presents SCOPER, an innovative tool that addresses these challenges by integrating deep-learning-based ion binding site prediction with conformational sampling, offering a more reliable approach to validate and refine RNA structures against experimental SAXS data. We provide our source code and a web server that runs the pipeline.

## Introduction

In recent years, novel and unexpected roles of non-coding RNAs have been discovered in multiple processes, such as signaling, cancer, development, and stress response (1, 2). Structural characterization of their solution conformations is critical to understanding their functional role (3). RNA flexibility challenges traditional structural characterization techniques, such as X-ray crystallography and NMR spectroscopy. While progress has been made with RNA structure determination by cryo-Electron Microscopy, challenges due to instability, heterogeneity, and small size limit widespread application (4). Consequently, RNA structures comprise only ∼3% of all structures in the Protein Data Bank (PDB) (5). Although novel deep learning methods produce highly accurate protein structures, for RNAs these models are not as accurate and can produce a wide variety of structural models with different base pairings (6).

Small-angle X-ray scattering (SAXS) can rapidly provide in-solution structural information on biological macromolecules, describing the size, shape, and dynamics (7–10). This in-solution structural technique is experiencing a revival primarily due to improvements in data collection technologies and computational algorithms (9). Unlike X-ray crystallography and NMR spectroscopy, SAXS is a fast and reliable technique performed under dilute conditions, thus requiring minimal amounts of RNA samples. The technique has provided reliable data on particles ranging from small RNA duplexes at 20 kDa to the large 70S ribosome at 2700 kDa (10–13). SAXS can be an invaluable tool for the structural biologist, supplementing the traditional high-resolution techniques; yet, the method has limitations for RNA that merit attention.

Directly comparing an RNA structure with a SAXS profile requires an accurate calculator of theoretical SAXS profiles from atomistic models. Current SAXS profile calculators do not accurately match the experimental SAXS RNA data within the noise (14). The difference between the theoretical and experimental SAXS profiles is attributed to the ion-induced changes in the hydration layer (15) and the conformational diversity of RNA (16, 17). The results from SAXS studies of counterion interactions with rigid RNA duplexes have greatly improved our understanding of highly charged RNA’s folding into compact structures (18). Ions, commonly Mg^2+^, also contribute signal to the SAXS profiles with the X-ray scattering length density of Mg^2+^ almost two times larger than H_2_O. However, the ions are not predicted by structure modeling algorithms and can be easily missed by experimental structure determination methods. In addition to the localization of ions, we need to be able to predict their solution conformations for functional studies of RNA molecules. Direct comparison between experimental or predicted RNA 3D structures and SAXS profiles measured in solution is challenging. A solution conformation can have similar secondary and tertiary structures as the structural model but different overall conformation due to the RNAs’ flexibility. Moreover, for larger flexible RNAs to describe conformational heterogeneity, multi-state models (two or more conformations and their weights) can be required (19, 20).

Novel works for directly predicting RNA structure from SAXS data have recently emerged such as Ernwin (21) and RNAMasonry (22). These works predict a tertiary structure beginning from a secondary structure and sample a coarse tertiary structure guided by SAXS data. However, these approaches do not account for Mg2+ ions or take multiple states into account.

Several methods exist to sample the conformational ensemble from a starting RNA structure, assuming the in-solution conformation only varies slightly. These include Molecular Dynamics simulations (19), Monte Carlo-based methods, such as SimRNA (23), Normal Modes (15, 24, 25), and robotics-inspired motion planning approach, KGSRNA (26). However, based on a statistical analysis of ions in the PDB, there is only one predictor of metal ion positions in RNA, MetalionRNA (27). MetalionRNA relies on a distance and angle-dependent anisotropic potential describing interactions between metal ions and RNA atom pairs. The statistical potential was calculated using ∼100 structures when it was first published. With the availability of novel geometric deep-learning methods for protein structures and a larger dataset of RNA structures, there is an opportunity to develop more accurate tools that predict metal ion positions. More recently, works leveraging large amounts of available data have approached this problem using deep learning, such as Zhou et al. (28), which uses 3D convolutional neural networks to comb the 3D structure of RNA as a 3D image. While this approach seems promising, 3D convolutions are not invariant to the arbitrary orientation of the RNA structure. 3D convolutions are also notoriously slow to compute and require large amounts of data to train while also dealing with many sparse voxels that hinder the training process. The newest version of Deepmind’s AlphaFold (AlphaFold3 (6)) released a powerful new model that can generate 3D RNA complexes along with predicted ion binding sites; however, their ion binding site prediction has no reported accuracy measures.

We developed a SAXS-based Conformation Predictor for RNA (SCOPER), which takes an initial RNA structure and a SAXS profile as input and outputs a single or multi-state model with improved fit to the SAXS profile and Mg^2+^ ions added to RNA conformations. We used geometric deep learning to train a model named IonNet for Mg^2+^ ion placement based on the atomic neighborhood. We use SAXS profiles to select subsets of ion positions that best fit experimental SAXS profiles to identify further the most probable ion positions from the predicted ones. To address the conformational flexibility of RNA molecules, we sample multiple conformations using KGSRNA, a motion planning algorithm that preserves the secondary structure of the initial structure, followed by the prediction of Mg^2+^ ion positions for each sampled conformation. Finally, we determine single or multi-state models that fit the data within the noise. KGSRNA sampling preserves the secondary structure of the initial RNA structure, enabling reliable sampling of plausible conformational changes driven by RNA flexibility in solution. By preserving RNA’s secondary structure, this approach minimizes the risk of overfitting SAXS data by models containing broken base pairs, which may happen in SAXS-guided normal mode sampling (29). We benchmarked SCOPER’s capability with 14 experimental SAXS datasets, including experimental data of three RNAs with known crystal structures (P4P6, SAM-riboswitch, and LYS-riboswitch) (30–32).

## Materials and Methods

### Summary

The input to our method is a PDB format structure of the RNA and a SAXS profile. The structure can be obtained experimentally or using RNA structure prediction tools, such as DeepRNAFold (33), RNAComposer (34), SimRNA (23), or recently released AlphaFold3 (6). The method proceeds in four main stages (Fig. 1). First, we generate 1,000 RNA conformations from the input structure using KGSRNA that perturbs the structure while preserving base pairing (26). The base pairing is assigned by RNAview (35) prior to the KGSRNA sampling. Second, IonNet predicts potential Mg^2+^ positions for each generated conformation. Third, the input SAXS profile selects a combination of Mg^2+^ ion positions that best fit the data using the goodness of fit parameter χ^2^. Finally, if a single conformation with Mg^2+^ ions does not fit the SAXS profile within the noise, we apply MultiFoXS to find multi-state models to explore conformational plasticity (36). If a SAXS profile is unavailable, it is still possible to use IonNet as a standalone tool, using the code on GitHub, to obtain the most probable Mg^2+^ ion positions using stages two and three only. Below, we describe stages two and three that we have developed specifically for this pipeline.

**Figure 1.**
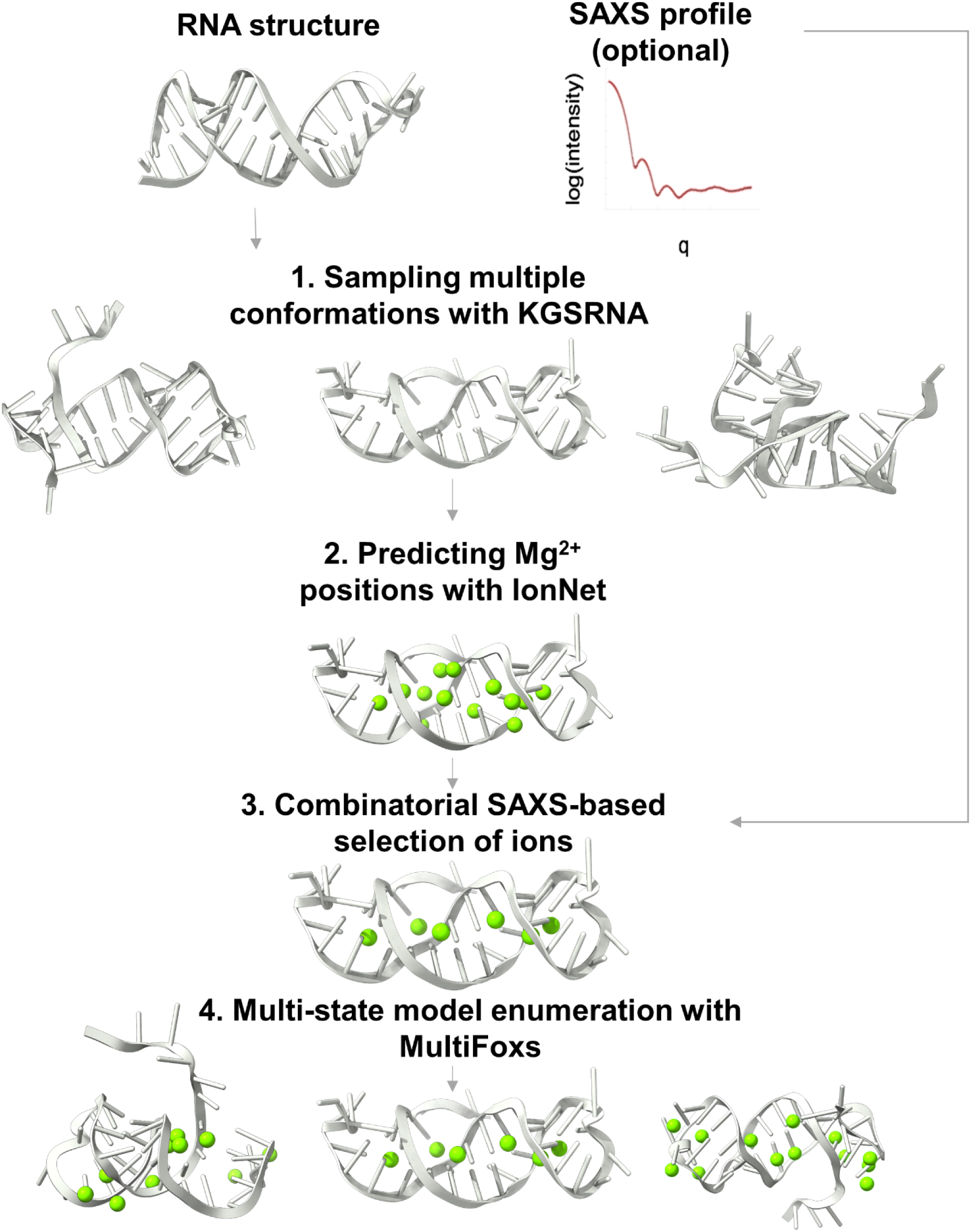
Visualization of SCOPER pipeline.

### Prediction of ion positions with IonNet

#### IonNet training

IonNet is a deep learning model that can classify a probe located on an RNA surface as either an Mg^2+^ ion or a water molecule based on the atomic neighborhood of the probe (atoms within an 8Å radius). To train the model, we relied on ∼1,000 PDB structures that contain ∼41,000 Mg^2+^ ions to serve as positive examples. Water molecules and their atomic neighborhoods were negative examples (Supplementary Data). We describe these neighborhoods as graphs, where nodes correspond to RNA atoms, and edges represent the distances between the nodes. Our model performs its classification using Graph Neural Networks (GNNs) leveraging graph attention (37) and graph convolution (38) layers to extract information from the input graphs (Fig. S1, Table S1). Four-fold cross-validation was performed with IonNet, resulting in an AUROC mean of 0.89 (Fig. 2A). We provide a complete overview of IonNet’s training and accuracy metrics in the Supplementary Data (Table S1, Table S3, Fig. S6-11).

**Figure 2:**
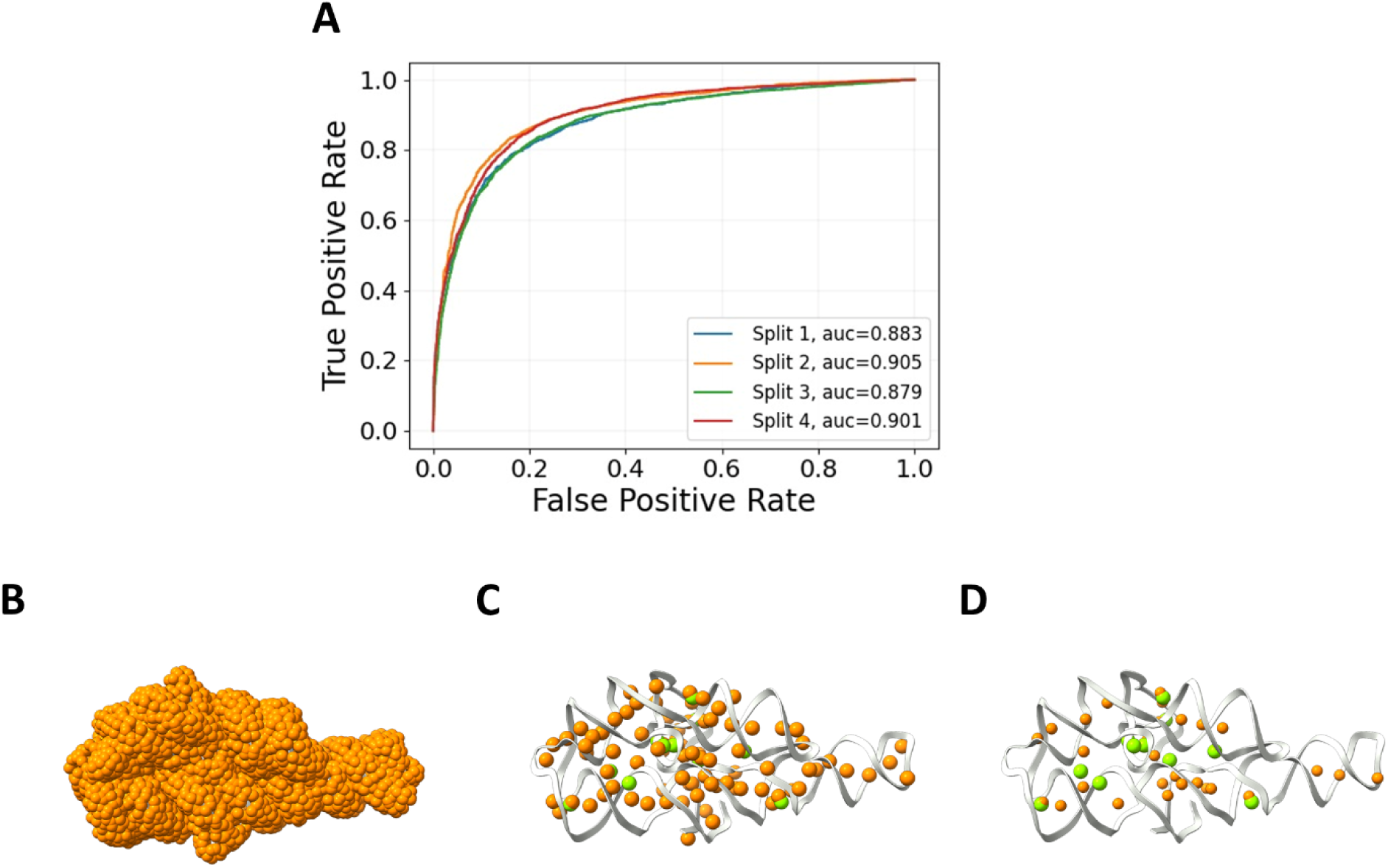
IonNet performance. **A.** Four-fold cross-validation of our best model over the whole test set. **B-D**. IonNet prediction of Mg^2+^ ions for the P4P6 structure (PDB 1GID). Ions are predicted from the set of 5,500 probes (orange) covering the whole structure (B) with high-confidence predictions above a threshold of 0.5 with ∼80 predictions (C) and 0.9 with ∼25 predictions (D). The experimentally observed Mg^2+^ ions are in green (12 in total).

#### IonNet inference

We generate surface probes to predict ion positions in a given RNA structure (Fig. 2B) using Connolly’s surface method (39). A neighborhood of RNA atoms (within 8Å) is extracted for each such probe. Each neighborhood is passed to IonNet and is classified as either an Mg^2+^ ion or a water molecule neighborhood. When IonNet is used outside of SCOPER as a standalone tool, iterative clustering selects the predicted binding sites with the highest confidence. By the end of this stage, IonNet suggests a number of plausible Mg^2+^ binding-site locations. We provide an accuracy metric for this inference process over our test set in the supplementary data.

For example, IonNet could identify probes in the vicinity of 11 out of 12 experimentally observed ions for the P4P6 structure (Fig. 2C). Overall, out of ∼5,500 probes, IonNet identified 84 as Mg^2+^ ions. If we use a strict cut-off (0.9 instead of 0.5) for Mg^2+^ ion prediction, 6 out of the 12 experimentally observed ions are correctly predicted, with 30 predicted Mg^2+^ ion positions (Fig. 2D).

### Combinatorial SAXS-based selection of ions

#### SAXS profile fitting

We computed the profiles using the Debye formula with the FoXS program (14, 40). For a given structural model, there are three adjustable parameters: excluded volume (*c_1_*), hydration layer density (*c_2_*), and the scaling factor (*c*) that are optimized to fit the experimental profile as measured by the χ^2^ score:

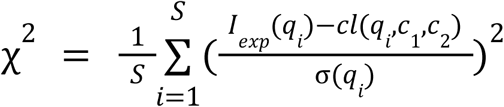

where *I_exp_(q)* is the experimental profile, which is a function of the momentum transfer *q* = 4π*sin*θ/λ, where 2θ is the scattering angle, λ is the wavelength of the incident X-ray beam, *σ(q)* is the experimental error of the measured profile, and *S* is the number of points in the profile. *I(q, c_1_, c_2_)* is the computed profile, given by the Debye formula (14), with *c_1_* and *c_2_* optimized to minimize the χ^2^ score.

### Enumeration of Mg^2+^ subsets

In the pipeline’s 3rd stage, the subset of Mg^2+^ ions that minimizes the χ^2^ score is selected from a set of predicted Mg^2+^ ion positions by IonNet. These subsets are enumerated using a branch-and-bound algorithm. First, we calculate the χ^2^ score for the RNA structure with a single ion. Second, we enumerate all possible subsets with two ions (a branch step) and keep *K* best scoring ones (a bound step) for the next iteration, when a third ion will be added. We continue enumerating the ion subsets of size *N* based on size *N − 1* subsets until the score can no longer be improved. To speed up the calculation of the SAXS profile and the corresponding χ^2^ score for each subset, we precompute the RNA profile and the profile of each Mg^2+^ ion relative to the RNA. This preprocessing enables a rapid summation of the relevant sub-profiles for each subset of Mg^2+^ ions and χ^2^ score calculation.

### SAXS data collection

To validate our pipeline, we collect experimental SAXS data from a monomeric RNA state free of contaminants (higher oligomeric states and aggregation). We applied Size Exclusion Chromatography coupled with small angle X-ray scattering (SEC-SAXS) (Tables S2, Fig. S2). All data, except RNA #10 and #14, were collected at SIBYLS beamline 12.3.1 at Advanced Light Source with recently reported developed SEC-SAXS-MALS modality (41). X-ray wavelength was set at λ=1.127 Å, and the sample to detector distance was 2100 mm, resulting in scattering vectors, *q*, ranging from 0.01 Å^-1^ to 0.4 Å^-1^. 60-95 μL of annealed RNAs with a concentration between 1 and 3 mg/ml were prepared in the SEC running buffer (Table S2). The Shodex KW802.5 column was equilibrated with a running buffer with a flow rate of 0.65 mL/min. Each sample was injected in an SEC column, and two-second X-ray exposures were recorded continuously for 24 min. RNA #10 and #14 were previously collected at the SIBYLS beamline and reported (13, 32). Program RAW (42) was used for further SEC-SAXS processing, including buffer subtractions and merging SAXS frames across the elution peak (Fig. S2). For RNA #8, #9, and #11, we applied the EFA approach (43) to ensure that the SAXS signal is derived from monomeric and well-folded RNA. The final merged SAXS curves were further used for Guinier analysis using the program RAW (42) and computing P(r) functions by the program GNOM (44). The MW_SAXS_ was calculated using Volume of Correlation (V_c_) (45) and compared to the molecular weight estimated by SEC-MALS (Table S2). The SEC-SAXS were deposited to the SIMPLE SCATTERING database https://simplescattering.com/ (deposition IDs are listed in Table S2).

## Results

### Impact of ionic strength and RNA plasticity on SAXS fitting

Fast calculation of SAXS profiles from protein structures usually involves modeling the hydration shell with implicit solvent models (14, 46–49). In these models, the density of the hydration layer can be adjusted to optimize the fit to the experimental SAXS profile (14, 46). The changes in the protein’s hydration layer, typically influenced by the protein surface net charge, result from the varying ionic strength of the buffer used in SAXS experiments (14). However, in the nucleic acid world, higher ionic strength increases the rigidity of nucleic acid structures (16) and can alter the overall arrangement of RNA segments (50, 51), whereas the hydration layer is poorly understood. The ionic strength-dependent plasticity of RNA poses a greater difficulty in quantifying changes in the RNA hydration layer using SAXS. Nevertheless, here, we look at the impact of salt concentration on the hydration layer model in the FoXS SAXS calculator (14) and RNA plasticity.

We measured SAXS profiles for a small RNA stem-loop (#3 in the benchmark) and P4P6 RNA (#13 in the benchmark) at different ionic strengths while keeping the Mg^2+^ concentration constant at 5 mM. Although the Radius of gyration (Rg) values obtained by the Guinier plot (Fig. 3A,B) are identical within the error across the ionic strengths (Table 1), we observed variations in the SAXS profile shown by the normalized Kratky plot (Fig. 3A,B). The narrowing of the peak at higher ionic strength (Fig. 3A,B) could be further visualized by changes in the P(r) function (Fig. 3C,D). Low concentration of RNA’s (stem-loop ∼0.06mM, P4P6 ∼0.01mM) at the elution peak, presence of 5mM Mg^2+^, and identical Rg’s across measured salt concentrations (Table 1) shows that SAXS-changes are not derived from altered inter-particular RNA interactions (52). Multiple factors may explain the observed changes: 1) changes in RNA rigidity as previously observed for DNA (53–55) and RNA rearrangement (50, 51), or changes in the hydration layer influenced by the altered RNA surface net charge, or all of the above. Nevertheless, because distinguishing RNA plasticity/rigidity from the altering hydration layer is difficult, we fit the SAXS profiles with the single or ensemble model of RNA, with/without adjustment of the c_1_/c_2_ parameters or with/without placement of Mg^2+^. The quality of fit for all the fittings is listed in Table 1. The single conformer with adjusted c_1_/c_2_ parameters did not match the SAXS profiles within the noise (Table 1), indicating that adding Mg^2+^ ions or a multi-state model (36) is necessary to match the data. However, the fit for a model with placed Mg^2+^ or a two-state model is similar (Table 1, Fig. 3E,F) and shows that the RNA plasticity or placement of Mg^2+^ is challenging to distinguish. The single conformer with placed Mg^2+^ ions and default c_1_/c_2_ parameters slightly worsened the fit. In contrast, multi-state models with placed Mg2+ ions and default c1/c2 parameters significantly improved the fit for P4P6 but not stem-loop (Table 1). This further indicates that fitting SAXS simultaneously with RNA plasticity, adjustment of the hydration layer and excluded volume, and placing Mg^2+^ may lead to data overfitting. To avoid SAXS data overfitting, we set the c1 and c*2* parameters to the default value of 1.0 in our further pipeline testing.

**Figure 3.**
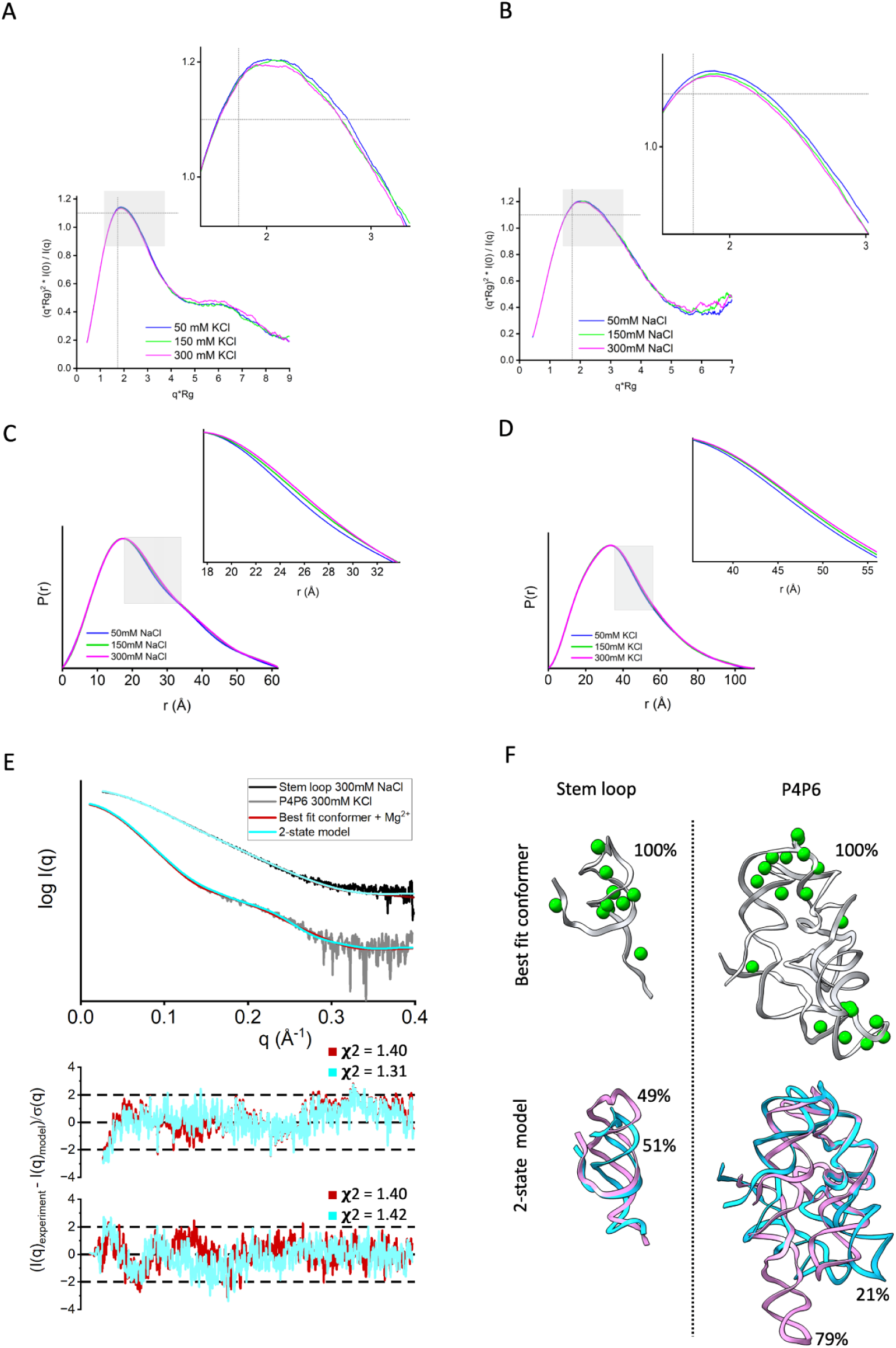
Impact of ionic strength on the hydration layer, Mg^2+^ placement, and RNA conformation. A-B Normalized Kratky plot for experimental SAXS curves of two RNAs (#3- RNA stem-loop, #13- P4P6) measured at three different salt concentrations, 50 mM (blue), 150 mM (green), and 300 mM (pink). The SAXS signal was smoothed using adjacent averaging to visualize differences between the curves better. The dashed gray lines on the Dimensionless (Rg) plot are guidelines for a globular protein. For a globular protein, the peak position should be at *qRg*=3≈1.73, while the peak height should be 3/*e*≈1.1. **B-C.** P(r) functions of two RNAs (#3- RNA stem-loop, #13- P4P6) calculated from SAXS curves (see Fig. S4) measured at three different salt concentrations, 50 mM (blue), 150 mM (green), and 300 mM (pink). **C.** Experimental SAXS curves collected at 300mM salt concentration for RNA #3 (stem-loop, black) and #13 (P4P6, gray). SAXS fit of the best-fit structure with Mg^2+^ (red) and two-state model without Mg^2+^ (cyan) reveal similar goodness of fit. The c_1_ and c_2_ parameters were adjusted in both fitting approaches (see Table 1). Bottom panel: SAXS fit residual with its ***χ***^2^ values. **D.** Best-fit conformer with Mg^2+^ and 2-state model shown for #3- RNA stem-loop, #13- P4P6 with corresponding weights. The ***χ***^2^ and c_2_ values for all salt concentrations are in Table 1.

**Table 1.**
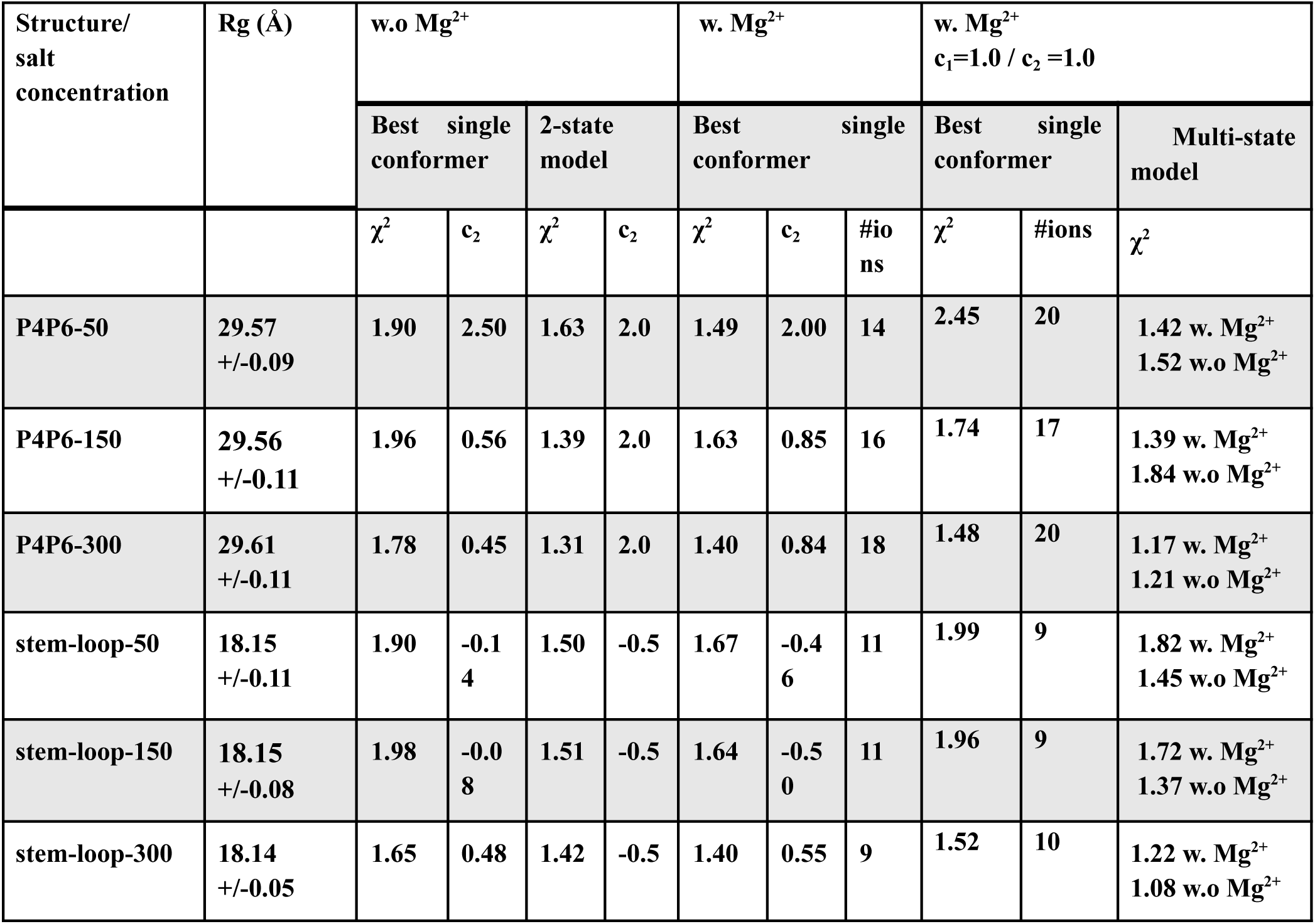
Concentration results.

| Structure/<br>salt<br>concentration | Rg (Å) | w.o Mg<sup>2+</sup> |  |  |  | w. Mg<sup>2+</sup> |  |  | w. Mg<sup>2+</sup><br>c<sub>1</sub>=1.0 / c<sub>2</sub>=1.0 |  |  |
| --- | --- | --- | --- | --- | --- | --- | --- | --- | --- | --- | --- |
|  |  | Best single<br>conformer |  | 2-state<br>model |  | Best single<br>conformer |  |  | Best single<br>conformer |  | Multi-state<br>model |
| | | $\chi^2$ | c<sub>2</sub> | $\chi^2$ | c<sub>2</sub> | $\chi^2$ | c<sub>2</sub> | #ions | $\chi^2$ | #ions | $\chi^2$ |
| P4P6-50 | 29.57<br>+/-0.09 | 1.90 | 2.50 | 1.63 | 2.0 | 1.49 | 2.00 | 14 | 2.45 | 20 | 1.42 w. Mg<sup>2+</sup><br>1.52 w.o Mg<sup>2+</sup> |
| P4P6-150 | 29.56<br>+/-0.11 | 1.96 | 0.56 | 1.39 | 2.0 | 1.63 | 0.85 | 16 | 1.74 | 17 | 1.39 w. Mg<sup>2+</sup><br>1.84 w.o Mg<sup>2+</sup> |
| P4P6-300 | 29.61<br>+/-0.11 | 1.78 | 0.45 | 1.31 | 2.0 | 1.40 | 0.84 | 18 | 1.48 | 20 | 1.17 w. Mg<sup>2+</sup><br>1.21 w.o Mg<sup>2+</sup> |
| stem-loop-50 | 18.15<br>+/-0.11 | 1.90 | -0.14 | 1.50 | -0.5 | 1.67 | -0.46 | 11 | 1.99 | 9 | 1.82 w. Mg<sup>2+</sup><br>1.45 w.o Mg<sup>2+</sup> |
| stem-loop-150 | 18.15<br>+/-0.08 | 1.98 | -0.08 | 1.51 | -0.5 | 1.64 | -0.50 | 11 | 1.96 | 9 | 1.72 w. Mg<sup>2+</sup><br>1.37 w.o Mg<sup>2+</sup> |
| stem-loop-300 | 18.14<br>+/-0.05 | 1.65 | 0.48 | 1.42 | -0.5 | 1.40 | 0.55 | 9 | 1.52 | 10 | 1.22 w. Mg<sup>2+</sup><br>1.08 w.o Mg<sup>2+</sup> |

### SAXS profile fitting

Our dataset consisted of 14 RNA samples with experimental SAXS profiles. The starting structures were obtained from DeepFoldRNA (#1-4, #6-8) (33), RNAComposer (#5, #9, #11), (34), and X-ray crystallography structures (#10 PDB 2GIS, #13, PDB 1GID, #14 PDB 3D0U) (31, 32, 56). We used different programs to predict our initial structures mainly due to time constraints, where RNAComposer produced initial fitting structures faster than DeepFoldRNA. DeepFoldRNA was used when the RNAComposer model fitted the SAXS data poorly. Additionally, we apply the SCOPER pipeline with AlphaFold3 for large RNAs (#9, #11) to show how our pipeline can validate or reject models with different secondary structures (see below). Deep learning prediction models such as DeepFoldRNA or AlphaFold3 offer multiple conformations when predicting 3D structures. We always selected the conformation with the best initial SAXS fit. We found that predictions with an inadequate initial structure were usually unsuitable for our pipeline as KGSRNA explores the conformational space of RNA by preserving its secondary structure. Our pipeline is expected to benefit from future improvements in RNA structure prediction that will provide more accurate starting structures (57). The fit of most starting structures to their corresponding SAXS profile was relatively poor (Table 2, Fig. 4). We predicted Mg^2+^ ion positions using IonNet. Adding Mg^2+^ ions to the starting structure only marginally improved the fit (Table 2, Fig. 4, Fig. 5). However, in the case of the accurately determined starting structure of the SAM-riboswitch (#10) obtained from X-ray crystallography, the addition of Mg^2+^ ions resulted in significant improvement (Table 2, Fig. 4). The SAM-riboswitch is a well-folded globular RNA where the addition of a relatively large number of placed Mg^2+^ ions contributes to the improvement of the SAXS profile fit. Significant improvement in SAXS fit could also be observed for some small RNAs (#3, #5) with more straightforward folds and expected limited plasticity. Thus, the starting predicted structure represents the solution state, and placing Mg^2+^ ions improves the SAXS fit. However, selecting a more accurate conformation is critical to matching the experimental SAXS profiles except for the RNA-riboswitch (#10).

**Figure 4.**
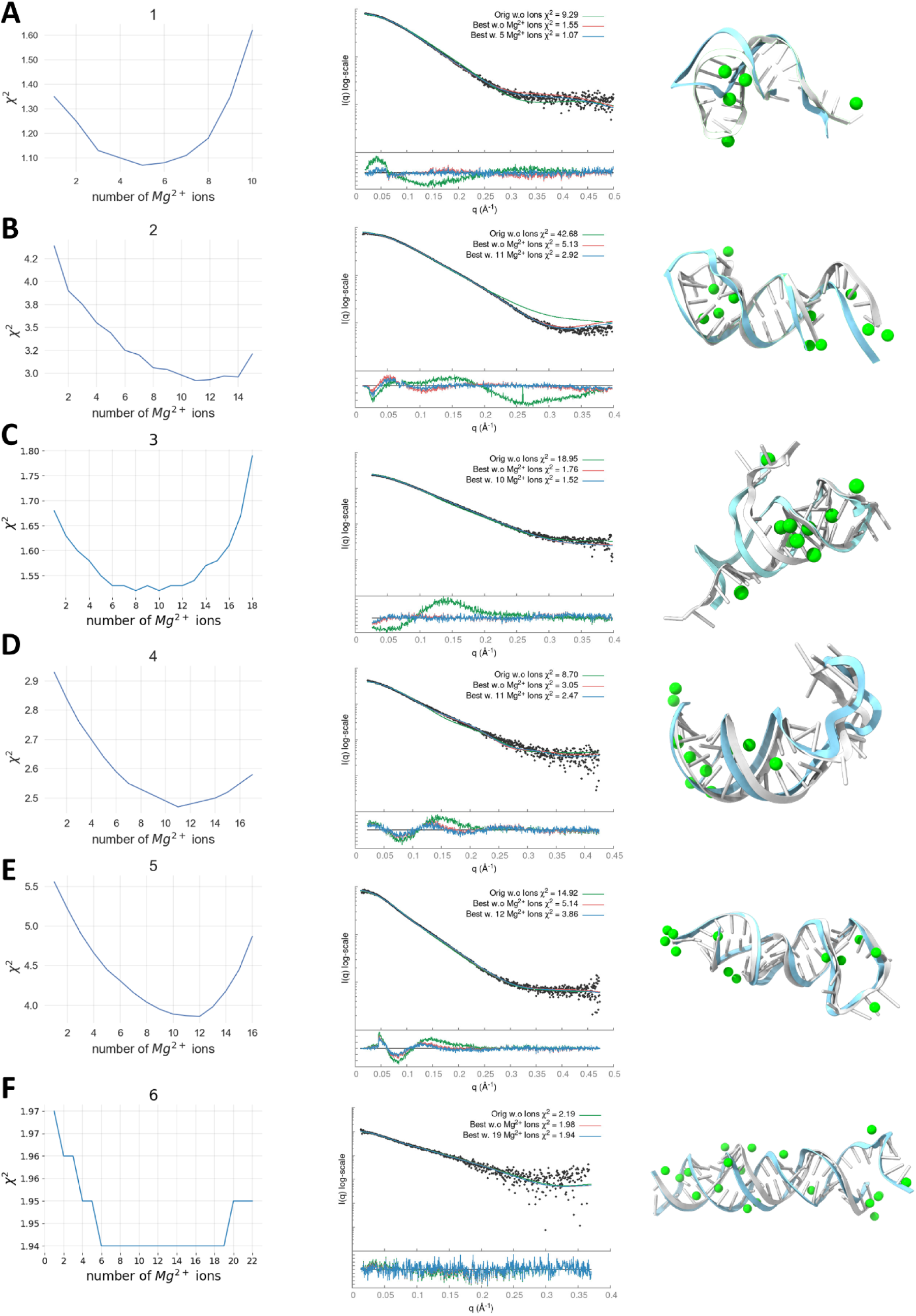
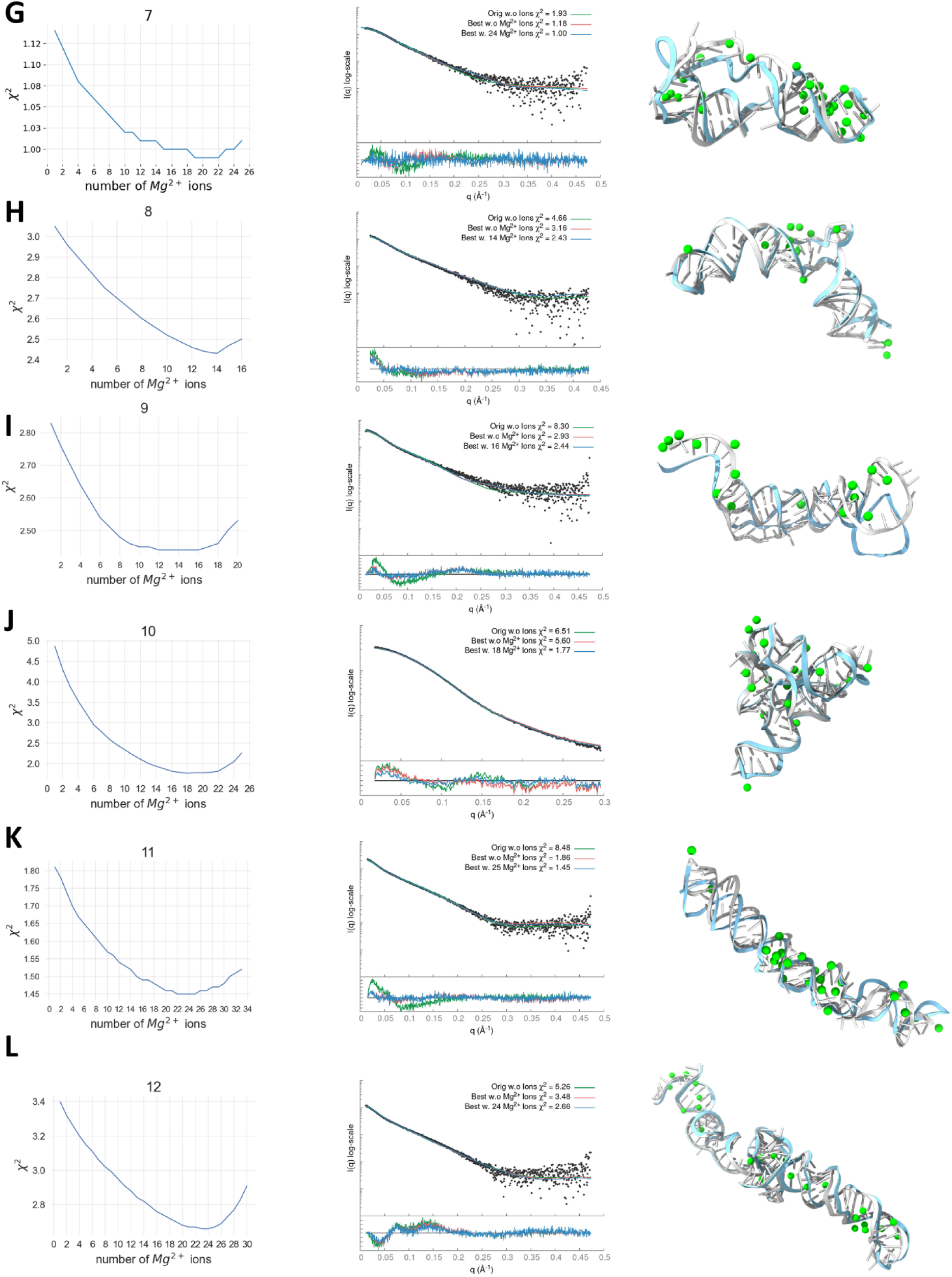
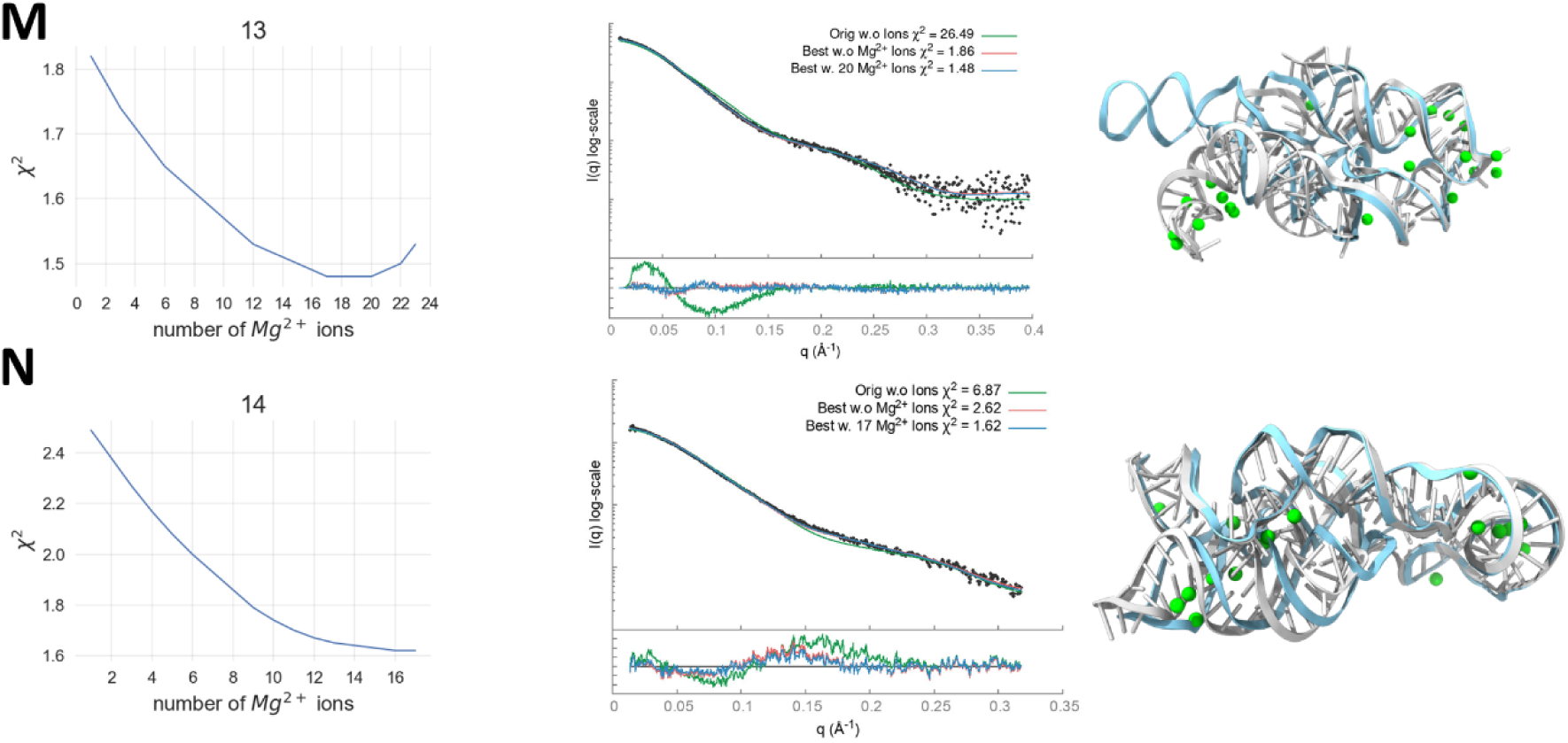
SCOPER results for 14 benchmark cases (labeled A to N). The decrease in χ^2^ per Mg^2+^ ion added (left column plots), SAXS profile fit of original starting structure (green), best structure without Mg^2+^ ions (red), and best structure with added Mg^2+^ ions (blue) vs. experimental profile (black dots) (middle column plots), initial RNA structure (cyan) and best fitting RNA structure (white) with predicted Mg^2+^ ions (green) (right column).

**Figure 5.**
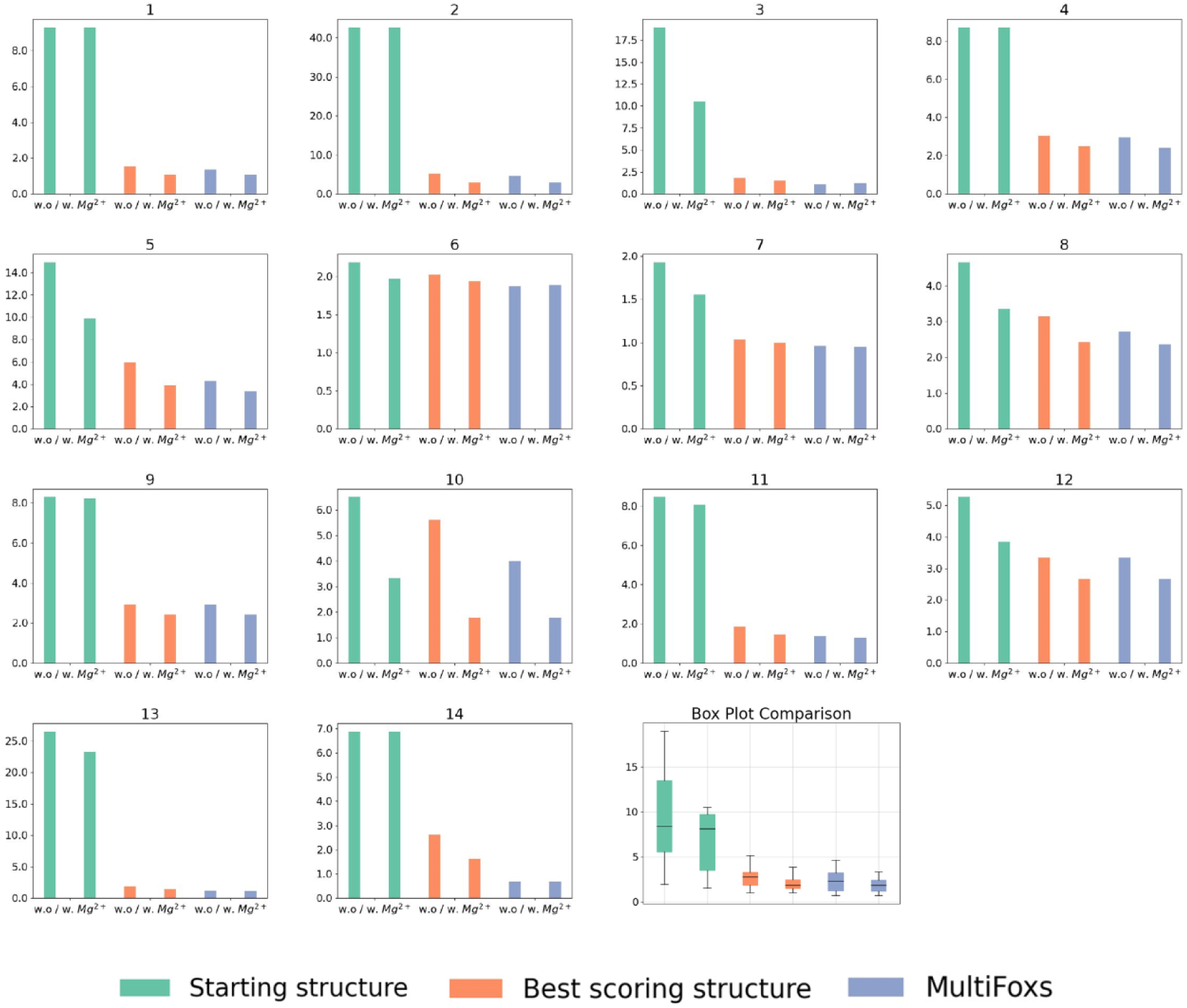
Bar plots of χ^2^ scores for the 14 benchmark cases: starting structures (green), best scoring conformations (orange), and multi-state models (purple), all with and without the addition of Mg^2+^ ions. A summary of the scores for all benchmark cases is provided in the last boxplot.

**Table 2.** *χ*^2^ values for our fitting experiments (corresponding to plots in Fig. 4.)

| RNA | Starting Structure |  | Best Scoring Structure |  | MultiFoXS |  | Number of States |
| --- | --- | --- | --- | --- | --- | --- | --- |
|  | w.o Mg<sup>2+</sup> | w. Mg<sup>2+</sup> | w.o Mg<sup>2+</sup> | w. Mg<sup>2+</sup> | w.o Mg<sup>2+</sup> | w. Mg<sup>2+</sup> |  |
| 1 | 9.29 | 9.29 | 1.55 | 1.07 | 1.35 | 1.06 | 3 |
| 2 | 42.67 | 42.67 | 5.12 | 2.92 | 4.64 | 2.92 | 3 |
| 3 | 18.95 | 10.52 | 1.76 | 1.52 | 1.08 | 1.22 | 6 |
| 4 | 8.70 | 8.70 | 3.04 | 2.47 | 2.94 | 2.41 | 3 |
| 5 | 14.91 | 9.90 | 5.95 | 3.86 | 4.27 | 3.35 | 3 |
| 6 | 2.18 | 1.97 | 2.02 | 1.94 | 1.87 | 1.89 | 3 |
| 7 | 1.92 | 1.55 | 1.03 | 1.00 | 0.96 | 0.95 | 6 |
| 8 | 4.65 | 3.35 | 3.15 | 2.43 | 2.71 | 2.37 | 4 |
| 9 | 8.30 | 8.20 | 2.93 | 2.44 | 2.93 | 2.44 | 2 |
| 10 | 6.51 | 3.33 | 5.60 | 1.77 | 4.00 | 1.77 | 2 |
| 11 | 8.48 | 8.07 | 1.86 | 1.45 | 1.37 | 1.31 | 4 |
| 12 | 5.26 | 3.84 | 3.34 | 2.66 | 3.34 | 2.66 | 2 |
| 13 | 26.49 | 23.24 | 1.86 | 1.48 | 1.21 | 1.17 | 4 |
| 14 | 6.87 | 6.87 | 2.62 | 1.62 | 0.69 | 0.69 | 4 |

In all of our benchmark cases, the single conformation with the best fit (lowest χ^2^) out of the 1,000 sampled by KGSRNA could fit the data significantly better than the initial structure (Table 2, Fig. 4,5). Adding Mg^2+^ ions to these conformations further improved the fit (Table 2, Fig. 4,5). Additional improvement in the fit to SAXS profiles could be achieved using multi-state models that mimic RNA plasticity (Table 2). Notably, the multi-segment RNAs where the more significant movement of segments could be expected (#11, #13, and #14) (50) showed significant fit improvement with a multi-state model. For example, the SAXS profile of the P4P6 had the best fit with four conformations (Fig. S3) with a χ^2^ =1.17 vs. 1.48 for a single conformation. However, the different placements of Mg^2+^ in each conformer (Fig. 3) suggest that placing Mg^2+^ ions in combination with multi-state modeling can lead to the overfitting of experimental SAXS data. On the other hand, it may also indicate that the movement of individual P4P6 segments in solution can result in more loosely defined coordinates for Mg^2+^ ions. Generally, validating through SAXS whether the Mg^2+^ ions observed in the P4P6 crystal structure remain ordered in the dynamic solution state is challenging. Smaller RNAs with single-stranded regions (#3) also show improved fit with a multi-state model. The multistate model of the #3 RNA mimics the plasticity of the unpaired 5’-end, resulting in an χ^2^ value of 1.22 compared to 1.52 for a single conformation (Table 2). These results indicate that our pipeline can predict Mg^2+^ binding sites that significantly reduce the χ^2^ score of the RNA structure (Figs. 4-5, Table 2). Although the improvements to the fit by adding Mg^2+^ ions were relatively marginal compared to finding a suitable conformation, these additions are not insignificant (Table 2, Figs. 4-5).

We conducted our profile fit calculations over all 1,000 conformations generated by KGSRNA. We found that running IonNet on all 1,000 conformations can be redundant if a single conformation fits the data within the noise. The best-fitting conformation without Mg^2+^ ions tends to be one of the best-fitting conformations with added ions (Fig. S5). If a user lacks parallel computing power, a safe assumption is that our pipeline can be run only on the best-scoring structures sampled by KGSRNA with little cost to the optimality of the output. However, if multi-state modeling is needed to fit the data within the noise, one needs to run IonNet for all the conformations.

### Using SCOPER for validation of structural models

We observed that the SCOPER can validate structures with adequate secondary and tertiary initial structures for the SAXS profile. We find that only adding Mg^2+^ ions predicted by IonNet is insufficient to fit the data (Table 2, Fig. S5) as most structures’ fit to the profile is usually only marginally changed when Mg^2+^ ions are added. IonNet selects binding sites with high accuracy, and SCOPER selects ions that reduce the SAXS score with a clustering algorithm; both help to reduce overfitting the SAXS profile by only adding ions, although these do not eliminate the possibility that some ions are chosen specifically because they help overfit the SAXS profile.

During experimentation, we also noticed that SCOPER worked best when the initial structure fit relatively well. Because KGSRNA preserves the initial structure’s secondary structure, SCOPER is unlikely to overfit a structure with the wrong secondary structure convincingly. While KGSRNA does lend itself to exploring a large conformation space (Fig. S12), we found 1,000 iterations to be sufficient for our needs; however, this number can be increased if needed.

We ran SCOPER with two initial models derived from two different prediction programs for three RNAs to demonstrate the role of SCOPER in validating structures rather than using it as a structure modeling tool. We chose two large RNAs (#9, #11) with the unknown experimentally defined structure and possible variations in tertiary structure prediction by AlphaFold3 (6) vs. DeepFoldRNA or RNAcomposer. The SCOPER pipeline shows a significantly better fit for the refined DeepFoldRNA models (#9 χ^2^ =3.65 vs. 2.44, #11 χ^2^ =5.84 vs. 1.45) (Fig. 6A, B). In case #9, the unfolded RNA 3’-end was the most significant difference contributing to the good fit derived from the DeepFoldRNA model. In case #11, the absence of unpaired stem-loop segments in the AlphaFold3 model leads to a noticeably poorer fit.

**Figure 6.**
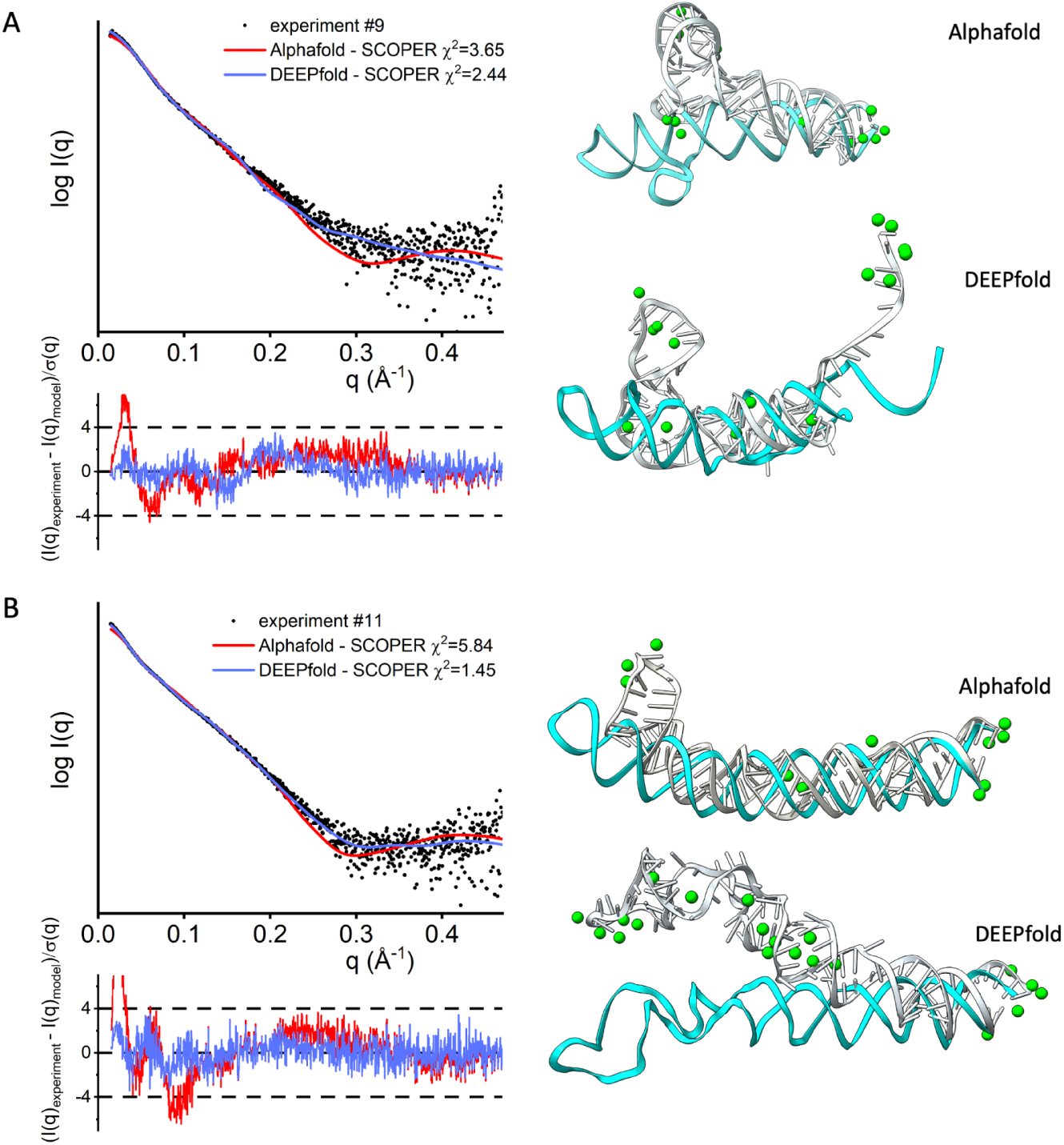
SCOPER results for benchmark cases #9 and #11 (labeled A and B). SAXS profile fit of AlphaFold3-refined structure (red), and DeepFoldRNA-refined structure with added Mg^2+^ ions (blue) vs. experimental profile (black dots), initial RNA structure (cyan) and best fitting RNA structure (gray) with predicted Mg^2+^ ions (green) (right column).

When multiple conformations exist in a solution with variations in secondary structure between them, SCOPER can be run with different initial structures. The resulting multi-state model can then be determined by combining SCOPER’s outputs for all initial structures using MultiFoXS (36). Therefore, we strongly recommend using SCOPER to validate or reject structural models based on a SAXS profile rather than relying on it solely as a predictive tool. This is because SCOPER’s effectiveness is greatly dependent on the accuracy of the initial structure, and its limitations must be carefully taken into account.

### Correlation between RNA size and a number of potential Mg^2+^ binding sites

We found a positive correlation (Pearson coefficient of 0.67) between the number of atoms in an RNA structure and the amount of predicted Mg^2+^ ions for the structure (Fig. 7). This is an expected outcome as the more surface area the structure has, it would seem likely that there would be more Mg^2+^ binding sites to stabilize the structure.

**Figure 7.**
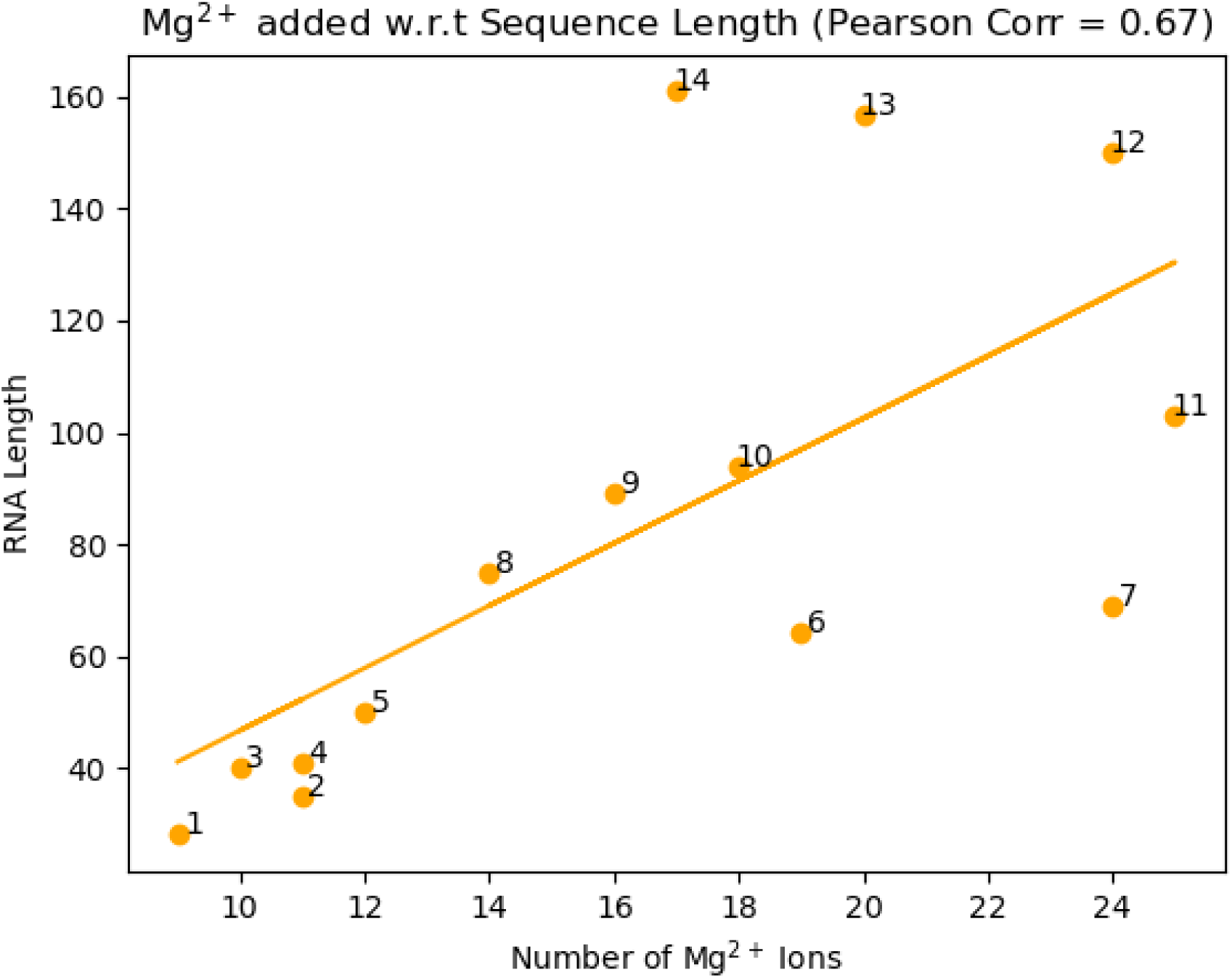
Number of Mg^2+^ ions (x-axis) vs. the sequence length (y-axis). The correlation coefficient is 0.670.

### SCOPER Webapp

We implemented a straightforward and easy-to-use web-based application to make the SCOPER pipeline available to the scientific community, particularly those without the knowledge or resources to install, configure, and run the Python code directly from the GitHub repository. The SCOPER pipeline has been added as an option within our existing BILBOMD job runner framework (58), available at https://bilbomd.bl1231.als.lbl.gov/. Users are required to create an account before submitting jobs. The required inputs are the initial RNA structure in PDB format and the experimental SAXS curve containing three columns (*q* in Å^-1^ unit, intensities, and experimental error.). The SCOPER web server implements the following pipeline steps (Fig. 1): (i) conformational sampling by KGSRNA, (ii) selection of the best conformer using FoXS, (iii) prediction of possible locations of Mg^2+^ ions using IonNet, and (iv) selection of the best Mg^2+^ ion placements by fitting the SAXS data using MultiFoXS. Users are notified by email when their jobs are complete. Results are then available to view directly within the web app (Fig. 8). They are also available to download as a compressed file containing the original uploaded RNA PDB file and the SAXS fit files, along with the output from the SCOPER pipeline.

**Figure 8.**
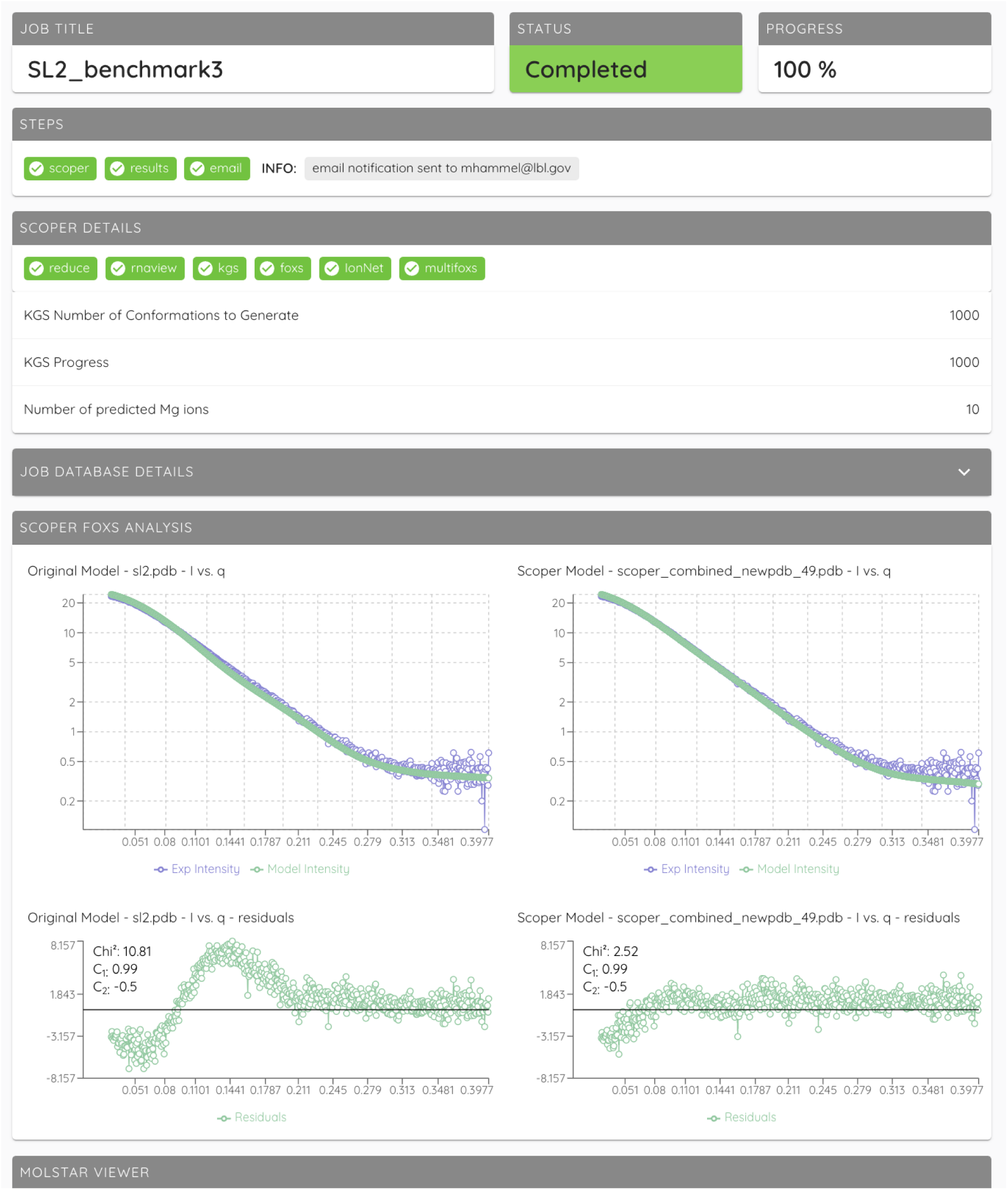
An example of the SCOPER job results page.

The SCOPER webapp executes its calculations with the single best initial conformer out of the KGSRNA samples. The user can choose to optimize the excluded volume, hydration layer, or adjustment of offset in the SAXS fitting parameters. However, we recommend that the calculations be fixed with the excluded volume and hydration layer parameters to 1 for initial modeling. MultiFoXS is not used in the webapp either, and only a single conformation with identified binding sites is returned as an output. We encourage users to report their results as such. More advanced users should be able to use our Github repository; however, they should be aware that by not setting the excluded volume and hydration layer parameters to 1 or using MultiFoXS, there’s a possibility of overfitting the data.

## Discussion and Conclusion

Although there has been substantial progress in predicting RNA structures, different tools can generate widely varying models for the same RNA molecule. In such cases, the experimental SAXS profile serves as a valuable resource for validating or discarding these models. For this purpose, we have developed SCOPER, a novel pipeline that integrates a deep learning model for placing Mg^2+^ ions into the RNA models or experimental structure, with SAXS-based validation to determine dynamic RNA conformation in solution. For ion placement, IonNet, a novel, deep learning-based model was trained to predict Mg^2+^ binding sites. Results show that these binding sites are predicted with high accuracy. Support for additional ions commonly found as structure stabilizers in the vicinity of RNA, such as Na^+^ and K^+,^ can be added with sufficient training data in future works.

The final ion positions and multi-state models are selected based on a fit to SAXS data. However, data overfitting is possible when predictions of ions used in conjunction with experimental SAXS profiles select a subset of Mg^2+^ ions that only improve the fit. To reduce overfitting, we recommend not varying adjustable SAXS fitting parameters, where adjusting the hydration layer may lead to the incorrect placement of Mg^2+^. However, with only three SAXS datasets (S-adenosylmethionine riboswitch #10, P4P6 #13, and lysine riboswitch #14) that have an experimentally verified structure, we cannot be sure that our pipeline entirely prevents overfitting. We suggest using a high threshold (0.3-0.5 for our model) to obtain only the most confident IonNet predictions.

Another potential concern is our dataset construction. Regarding the accuracy of each Mg²⁺ ion position, errors commonly arise because they are isoelectronic with water and Na⁺, leading to potential misidentification (59). By taking high resolution structures, we attempt to mitigate this issue. We also leverage neural networks’s ability to learn with noisy labels with large enough amounts of data (60) (61). We assume our model could learn despite these possibly noisy labels due to the model having a high precision and low recall. This is because a low recall value but a high precision may indicate that the model could learn true binding motifs while false positive labels would have no such discernable pattern. IonNet can be improved further by explicitly considering Mg^2+^ coordination when classifying binding sites. Most Mg^2+^ ions are generally coordinated by oxygen and nitrogen atoms (62). In future work, we plan an additional data post-processing step that classifies the coordination neighborhood based on distances to oxygen and nitrogen atoms. This classification can help improve the model’s precision even further.

We show that an accurate RNA SAXS calculator relies on the presence of Mg^2+^ ions in the model. The often-used extreme adjustment of the hydration layer, in the absence of Mg^2+^ ions, can lead to the wrong selection of RNA conformations by SAXS fitting. Optimizing the predicted starting RNA structural model is crucial in delivering a good agreement with experimental SAXS profiles. Although the prediction of Mg^2+^ ions in the RNA structure delivers a secondary improvement in the SAXS fit, it increases confidence in the RNA structure validation. This is particularly important when novel RNA structure prediction tools, like AlphaFold3 (26), need to be validated. We also note that SCOPER is limited to validating or discarding structures and is highly dependent on the accuracy of the initial structural model.

As shown in our comparison of χ^2^ for all models sampled by KGSRNA with and without Mg^2+^ (Fig. S5), adding the ions to the wrong model does not fit data better than the best conformer without Mg^2+^. Also, using different folds of the same RNA (Fig. 6) shows that adding Mg^2+^ to the wrong structure doesn’t fit data better than more correct predictions. However, the accuracy of the modeling, whether it is selecting the conformer or placing the Mg^2+^, also depends on data quality and the amount of RNA flexibility. Therefore the SCOPER pipeline should be considered a validation rather than a structure-determination tool in the relationship to SAXS data quality.

Overall, SCOPER’s results prove that it can help find more accurate RNA structures and suggest their conformations that better represent the solution state. The fact that the multistate model did not significantly improve the SAXS fit in most cases suggests that accounting for the Mg^2+^ in the RNA model is important to fit the experimental SAXS data properly. Here, we show that precisely calculated theoretical SAXS profiles can be used to validate or discard RNA structure predictions.

## Supporting information

Supplemental Paper

## Data Availability

The SEC-SAXS were deposited to the SIMPLE SCATTERING database (simplescattering.com). Additionally, the depositions contain the final merged SAXS curve and the final atomistic models used to calculate the SAXS fit (see Table S2). The RNA samples #4, #5, #7, #8, #9, #11, and #12 were collected under a proprietary agreement and unavailable in the SIMPLE SCATTERING database. The method is available from https://github.com/dina-lab3d/IonNet. For the latest release, please use our Zenodo link <u>dina-lab3D/IonNet: Zenodo Release</u> A SCOPER pipeline is available as a web server from https://bilbomd.bl1231.als.lbl.gov/.

## Author Contributions

D.S. and M.H. designed the research. Data curation and SAXS experiments were performed by M.H. The methodology, software development, and benchmarking were carried out by E.P. Data analysis, visualization, and writing of the manuscript were performed by E.P., D.S., and M.H. Writing of the paper was done by E.P, D.S., and M.H. S.C. contributed to data analysis and web-app development.

## References

1. Mattick, J.S., and I.V. Makunin. 2006. Non-coding RNA. Human Molecular Genetics. 15:R17–R29.

2. Esteller, M. 2011. Non-coding RNAs in human disease. Nature Reviews Genetics. 12:861–874.

3. Novikova, I.V., S.P. Hennelly, C.-S. Tung, and K.Y. Sanbonmatsu. 2013. Rise of the RNA Machines: Exploring the Structure of Long Non-Coding RNAs. Journal of Molecular Biology. 425:3731–3746.

4. Ma, H., X. Jia, K. Zhang, and Z. Su. 2022. Cryo-EM advances in RNA structure determination. Signal Transduct Target Ther. 7:58.

5. Berman, H.M. 2000. The Protein Data Bank. Nucleic Acids Research. 28:235–242.

6. Abramson, J., J. Adler, J. Dunger, R. Evans, T. Green, A. Pritzel, O. Ronneberger, L. Willmore, A.J. Ballard, J. Bambrick, S.W. Bodenstein, D.A. Evans, C.-C. Hung, M. O’Neill, D. Reiman, K. Tunyasuvunakool, Z. Wu, A. Žemgulytė, E. Arvaniti, C. Beattie, O. Bertolli, A. Bridgland, A. Cherepanov, M. Congreve, A.I. Cowen-Rivers, A. Cowie, M. Figurnov, F.B. Fuchs, H. Gladman, R. Jain, Y.A. Khan, C.M.R. Low, K. Perlin, A. Potapenko, P. Savy, S. Singh, A. Stecula, A. Thillaisundaram, C. Tong, S. Yakneen, E.D. Zhong, M. Zielinski, A. Žídek, V. Bapst, P. Kohli, M. Jaderberg, D. Hassabis, and J.M. Jumper. 2024. Accurate structure prediction of biomolecular interactions with AlphaFold 3. Nature.

7. Crist, B. 1983. A review of: *“SMALL ANGLE X-RAY SCATTERING”*, edited by O. Glatter and O. Kratky (Universitat Graz, Austria) New York: Academic Press, 1982, 515 pp. $89.50. ISBN 0-12-286280. Chemical Engineering Communications. 22:377–378.

8. Feigin, L.A., and D.I. Svergun. 1987. Structure Analysis by Small-Angle X-Ray and Neutron Scattering. .

9. Putnam, C.D., M. Hammel, G.L. Hura, and J.A. Tainer. 2007. X-ray solution scattering (SAXS) combined with crystallography and computation: defining accurate macromolecular structures, conformations and assemblies in solution. Quarterly Reviews of Biophysics. 40:191–285.

10. Hammel, M. 2012. Validation of macromolecular flexibility in solution by small-angle X-ray scattering (SAXS). European Biophysics Journal. 41:789–799.

11. Svergun, D.I., N. Burkhardt, J. Skov Pedersen, M.H.J. Koch, V.V. Volkov, M.B. Kozin, W. Meerwink, H.B. Stuhrmann, G. Diedrich, and K.H. Nierhaus. 1997. Solution scattering structural analysis of the 70 s Escherichia coli ribosome by contrast variation. II †. A model of the ribosome and its RNA at 3.5 nm resolution 1 †Paper I in this series is the accompanying paper, Svergun et al. (1997) 1Edited by M. F. Moody. Journal of Molecular Biology. 271:602–618.

12. Lipfert, J., and S. Doniach. 2007. Small-Angle X-Ray Scattering from RNA, Proteins, and Protein Complexes. Annual Review of Biophysics and Biomolecular Structure. 36:307–327.

13. Rambo, R.P., and J.A. Tainer. 2010. Improving small-angle X-ray scattering data for structural analyses of the RNA world. RNA. 16:638–646.

14. Schneidman-Duhovny, D., M. Hammel, J.A. Tainer, and A. Sali. 2013. Accurate SAXS profile computation and its assessment by contrast variation experiments. Biophys. J. 105:962–974.

15. Kirmizialtin, S., S.A. Pabit, S.P. Meisburger, L. Pollack, and R. Elber. 2012. RNA and Its Ionic Cloud: Solution Scattering Experiments and Atomically Detailed Simulations. Biophysical Journal. 102:819–828.

16. Chen, H., S.P. Meisburger, S.A. Pabit, J.L. Sutton, W.W. Webb, and L. Pollack. 2012. Ionic strength-dependent persistence lengths of single-stranded RNA and DNA. Proceedings of the National Academy of Sciences. 109:799–804.

17. Drozdetski, A.V., I.S. Tolokh, L. Pollack, N. Baker, and A.V. Onufriev. 2016. Opposing Effects of Multivalent Ions on the Flexibility of DNA and RNA. Physical Review Letters. 117.

18. Chen, Y., and L. Pollack. 2016. SAXS studies of RNA: structures, dynamics, and interactions with partners. Wiley Interdiscip. Rev. RNA. 7:512–526.

19. Chen, Y.-L., T. Lee, R. Elber, and L. Pollack. 2019. Conformations of an RNA Helix-Junction-Helix Construct Revealed by SAXS Refinement of MD Simulations. Biophys. J. 116:19–30.

20. Dethoff, E.A., J. Chugh, A.M. Mustoe, and H.M. Al-Hashimi. 2012. Functional complexity and regulation through RNA dynamics. Nature. 482:322–330.

21. Thiel, B.C., G. Bussi, S. Poblete, and I.L. Hofacker. 2024. Sampling globally and locally correct RNA 3D structures using Ernwin, SPQR and experimental SAXS data. Nucleic Acids Res. 52:e73.

22. Chojnowski, G., R. Zaborowski, M. Magnus, S. Mukherjee, and J.M. Bujnicki. 2023. RNA 3D structure modeling by fragment assembly with small-angle X-ray scattering restraints. Bioinformatics. 39.

23. Boniecki, M.J., G. Lach, W.K. Dawson, K. Tomala, P. Lukasz, T. Soltysinski, K.M. Rother, and J.M. Bujnicki. 2016. SimRNA: a coarse-grained method for RNA folding simulations and 3D structure prediction. Nucleic Acids Res. 44:e63.

24. Zacharias, M., and H. Sklenar. 2000. Conformational deformability of RNA: a harmonic mode analysis. Biophys. J. 78:2528–2542.

25. Sabei, A., T.G.C. Baia, R. Saffar, J. Martin, and E. Frezza. Internal Normal Mode Analysis applied to RNA flexibility and conformational changes. .

26. Fonseca, R., H. van den Bedem, and J. Bernauer. 2015. KGSrna: Efficient 3D Kinematics-Based Sampling for Nucleic Acids. Lecture Notes in Computer Science. 80–95.

27. Philips, A., K. Milanowska, G. Lach, M. Boniecki, K. Rother, and J.M. Bujnicki. 2012. MetalionRNA: computational predictor of metal-binding sites in RNA structures. Bioinformatics. 28:198–205.

28. Zhou, Y., and S.-J. Chen. 2022. Graph deep learning locates magnesium ions in RNA. QRB Discov. 3.

29. Manalastas-Cantos, K., P.V. Konarev, N.R. Hajizadeh, A.G. Kikhney, M.V. Petoukhov, D.S. Molodenskiy, A. Panjkovich, H.D.T. Mertens, A. Gruzinov, C. Borges, C.M. Jeffries, D.I. Svergun, and D. Franke. 2021. ATSAS 3.0: expanded functionality and new tools for small-angle scattering data analysis. J. Appl. Crystallogr. 54:343–355.

30. Battle, D.J., and J.A. Doudna. 2002. Specificity of RNA-RNA helix recognition. Proc. Natl. Acad. Sci. U. S. A. 99:11676–11681.

31. Montange, R.K., and R.T. Batey. 2006. Structure of the S-adenosylmethionine riboswitch regulatory mRNA element. Nature. 441:1172–1175.

32. Garst, A.D., A. Héroux, R.P. Rambo, and R.T. Batey. 2008. Crystal structure of the lysine riboswitch regulatory mRNA element. J. Biol. Chem. 283:22347–22351.

33. Pearce, R., G.S. Omenn, and Y. Zhang. *De Novo* RNA Tertiary Structure Prediction at Atomic Resolution Using Geometric Potentials from Deep Learning. .

34. Biesiada, M., K.J. Purzycka, M. Szachniuk, J. Blazewicz, and R.W. Adamiak. 2016. Automated RNA 3D Structure Prediction with RNAComposer. Methods Mol. Biol. 1490:199–215.

35. Yang, H., F. Jossinet, N. Leontis, L. Chen, J. Westbrook, H. Berman, and E. Westhof. 2003. Tools for the automatic identification and classification of RNA base pairs. Nucleic Acids Res. 31:3450–3460.

36. Schneidman-Duhovny, D., M. Hammel, J.A. Tainer, and A. Sali. 2016. FoXS, FoXSDock and MultiFoXS: Single-state and multi-state structural modeling of proteins and their complexes based on SAXS profiles. Nucleic Acids Research. 44:W424–W429.

37. Veličković, P., G. Cucurull, A. Casanova, A. Romero, P. Liò, and Y. Bengio. 2017. Graph Attention Networks. .

38. Rossi, E., F. Monti, M. Bronstein, and P. Liò. 2019. ncRNA Classification with Graph Convolutional Networks. .

39. Connolly, M.L. 1983. Analytical molecular surface calculation. Journal of Applied Crystallography. 16:548–558.

40. Schneidman-Duhovny, D., M. Hammel, and A. Sali. 2010. FoXS: a web server for rapid computation and fitting of SAXS profiles. Nucleic Acids Research. 38:W540–W544.

41. Rosenberg, D.J., G.L. Hura, and M. Hammel. 2022. Size exclusion chromatography coupled small angle X-ray scattering with tandem multiangle light scattering at the SIBYLS beamline. Methods Enzymol. 677:191–219.

42. Hopkins, J.B., R.E. Gillilan, and S. Skou. 2017. *BioXTAS RAW*: improvements to a free open-source program for small-angle X-ray scattering data reduction and analysis. Journal of Applied Crystallography. 50:1545–1553.

43. Meisburger, S.P., A.B. Taylor, C.A. Khan, S. Zhang, P.F. Fitzpatrick, and N. Ando. 2016. Domain Movements upon Activation of Phenylalanine Hydroxylase Characterized by Crystallography and Chromatography-Coupled Small-Angle X-ray Scattering. J. Am. Chem. Soc. 138:6506–6516.

44. Semenyuk, A.V., and D.I. Svergun. 1991. GNOM – a program package for small-angle scattering data processing. Journal of Applied Crystallography. 24:537–540.

45. Rambo, R.P., and J.A. Tainer. 2013. Accurate assessment of mass, models and resolution by small-angle scattering. Nature. 496:477–481.

46. Svergun, D., C. Barberato, and M.H.J. Koch. 1995. CRYSOL– a program to evaluate X-ray solution scattering of biological macromolecules from atomic coordinates. J. Appl. Crystallogr. 28:768–773.

47. Liu, H., R.J. Morris, A. Hexemer, S. Grandison, and P.H. Zwart. 2012. Computation of small-angle scattering profiles with three-dimensional Zernike polynomials. Acta Crystallogr. A. 68:278–285.

48. Poitevin, F., H. Orland, S. Doniach, P. Koehl, and M. Delarue. 2011. AquaSAXS: a web server for computation and fitting of SAXS profiles with non-uniformally hydrated atomic models. Nucleic Acids Res. 39:W184–9.

49. Virtanen, J.J., L. Makowski, T.R. Sosnick, and K.F. Freed. 2011. Modeling the hydration layer around proteins: applications to small- and wide-angle x-ray scattering. Biophys. J. 101:2061–2069.

50. Bai, Y., V.B. Chu, J. Lipfert, V.S. Pande, D. Herschlag, and S. Doniach. 2008. Critical assessment of nucleic acid electrostatics via experimental and computational investigation of an unfolded state ensemble. J. Am. Chem. Soc. 130:12334–12341.

51. Bruetzel, L.K., T. Gerling, S.M. Sedlak, P.U. Walker, W. Zheng, H. Dietz, and J. Lipfert. 2016. Conformational Changes and Flexibility of DNA Devices Observed by Small-Angle X-ray Scattering. Nano Lett. 16:4871–4879.

52. Qiu, X., K. Andresen, L.W. Kwok, J.S. Lamb, H.Y. Park, and L. Pollack. 2007. Inter-DNA attraction mediated by divalent counterions. Phys. Rev. Lett. 99:038104.

53. Geller, K., and K.E. Reinert. 1980. Evidence for an increase of DNA contour length at low ionic strength. Nucleic Acids Res. 8:2807–2822.

54. Shliakhtenko, L.S., I.L. Liubchenko, B.K. Chernov, and V.B. Zhurkin. 1990. [The effect of temperature and ionic strength on the electrophoretic motility of synthetic DNA fragments]. Mol. Biol. . 24:79–95.

55. 1999. DNA structure: cations in charge? Curr. Opin. Struct. Biol. 9:298–304.

56. Cate, J.H., A.R. Gooding, E. Podell, K. Zhou, B.L. Golden, C.E. Kundrot, T.R. Cech, and J.A. Doudna. 1996. CRYSTAL STRUCTURE OF A GROUP I RIBOZYME DOMAIN: PRINCIPLES OF RNA PACKING. .

57. Das, R., R.C. Kretsch, A.J. Simpkin, T. Mulvaney, P. Pham, R. Rangan, F. Bu, R.M. Keegan, M. Topf, D.J. Rigden, Z. Miao, and E. Westhof. 2023. Assessment of three-dimensional RNA structure prediction in CASP15. Proteins. 91:1747–1770.

58. Pelikan, M., G.L. Hura, and M. Hammel. 2009. Structure and flexibility within proteins as identified through small angle X-ray scattering. Gen. Physiol. Biophys. 28:174–189.

59. Leonarski, F., L. D’Ascenzo, and P. Auffinger. 2017. Mg2+ ions: do they bind to nucleobase nitrogens? Nucleic Acids Res. 45:987–1004.

60. Rolnick, D., A. Veit, S. Belongie, and N. Shavit. 2017. Deep learning is robust to massive label noise. arXiv [cs.LG].

61. Song, H., M. Kim, D. Park, Y. Shin, and J.-G. Lee. 2023. Learning From Noisy Labels With Deep Neural Networks: A Survey. IEEE Trans Neural Netw Learn Syst. 34:8135–8153.

62. Zheng, H., I.G. Shabalin, K.B. Handing, J.M. Bujnicki, and W. Minor. 2015. Magnesium-binding architectures in RNA crystal structures: validation, binding preferences, classification and motif detection. Nucleic Acids Res. 43:3789–3801.

