## Supplemental Paper for "Predicting RNA Structure and Dynamics with Deep Learning and Solution Scattering"

### Supplementary Material

#### 1. IonNet Training and Metrics.

**Data generation.** We downloaded 1,407 PDB format files from the Protein Data Bank (PDB) (1) that contain RNA and $Mg^{2+}$ atoms with a resolution below 3Å. We extracted all $Mg^{2+}$ ions within 8Å from the RNA molecule for each structure. These ions serve as our positive samples. We also randomly selected water molecules within 8Å from the RNA molecule to serve as our negative samples. We selected samples with a ratio of 1.5 water molecules per one $Mg^{2+}$ ion. In total, 41,725 $Mg^{2+}$ ions and 65,482 water molecules were selected. We augmented this dataset by generating additional $Mg^{2+}$ and water positions with the purpose of reducing sensitivity to the accuracy of the positions during the inference stage. To generate these positions, we relied on the probe centers of the solvent-accessible surface (2). The probe was placed at 1.4Å from the RNA surface, which we found to be the average distance for most $Mg^{2+}$ ions from the RNA in our dataset (Fig. S6). The density of the probes was set to 1 probe per $Å^2$ to optimize the trade-off between the accuracy of the probe position and the runtime performance of the inference stage. Probes were labeled as positive if they were at a distance of 3Å from an existing $Mg^{2+}$ ion and negative otherwise. In total 273,900 such probes were added to the dataset, with ~90,000 positive samples and the rest being negative samples.

**Data separation into train, validation, and test sets.** To construct non-overlapping train, validation, and test sets, we ran an exhaustive pairwise sequence alignment between all the RNA chains in our dataset. The dataset construction consisted of three main stages. First, we removed all sequences that had high sequence identity to many other sequences. Second, we detected similar pairs of sequences to prevent them from being separated into the train and test sets. Third, we divided the remaining sequences into the train, test, and validation sets in accordance with the marked pairs from the second stage.

In the first stage, we calculated an average sequence identity to all other sequences. This enabled us to discard very short or very long sequences that had high sequence identity to a large fraction of other sequences in the dataset. A sequence was discarded if the average sequence identity was above 56%, resulting in a more divergent selection of sequences, as seen in the similarity matrix (Fig. S7.). This filtering resulted in 961 sequences down from the original 1,538 RNA sequences. Measuring sequence identity scores between pairs of sequences with significantly different lengths results in different score distributions compared to score distributions measured between sequences of similar lengths. For this reason, we needed to obtain the distribution of sequence identity scores between sequences of differing lengths.

To solve this problem, in the second stage, all remaining sequences were placed into bins in accordance with their length. Small-sized bins were merged together with larger bins of similar lengths. We ended up with nine bins with a similar number of RNA chains (Table S3). Similar pairs were banned from being separated into the train and test sets if either their sequence identity was above 90% or if their sequence identity scores were in the top 1% in the score distribution between all pairs in both bins.

In the third stage, we randomly add 5% of the 961 filtered sequences into our test set. All the chains derived from that PDB structure are also included in the test set. For each of the remaining 95% of sequences we compared each potential chain to the RNA sequences in the test set. If the sequence was not banned from the training set for being too similar to a sequence in the test set, then it was added to the training set. Otherwise, it was added to the validation set. In total our best performing model used 580 chains for training, 91 for validation and 47 for testing. These come from 499 PDB files, 447 solved with X-ray, 49 with EM and 3 with NMR. This corresponds to 19,855 $Mg^{2+}$ ion positions for training, 3324 for validation and 638 for testing. With negative samples and augmented data included, train data used 45,753 positive samples and 113,941 negative samples, validation used 5770 positive and 18,628 negatives and the test set consisted of 1371 positives and 3584 negatives. To our knowledge these are the largest training and test sets used to train and evaluate models of a similar nature to IonNet.

**Problem definition.** The goal of our model is to take in as input an embedding of a neighborhood of either $Mg^{2+}$, water molecules, or surface probes near RNA atoms. We define the neighborhood of the ion, water molecule or probe as all atoms in a 8Å radius. The model then must label the neighborhood embedding of $Mg^{2+}$ or probes near (up to 3Å) to such an ion as positive and neighborhoods of water or probes far away (further than 3Å) from $Mg^{2+}$ ions as negative. For our models, we experimented with two main architectures: 3D-Convolutional Neural Networks (3D CNNs) and Graph Neural Networks (GNNs). Our embedding functions either embed neighborhoods into a 3D voxelized grid or into a graph where every node represents a single atom. The convolutional models proved ineffective, so in this work, we only discuss the training process and metrics for the GNNs and compare the results to the 3D CNN architectures.

**Feature definition.** Each node in the graph received a vector embedding of the physicochemical properties of each RNA atom. These atoms are characterized using the following 12 atom types according to SYBYL mol2 definition: carbon (C.2, C.3 and C.ar), nitrogen (N.2, N.4, N.am, N.ar, and N.p13), oxygen (O.2, O.3 and O.co2), phosphate (P.3) and sulfur (S.3). Additional features include the solvent accessible area of the atom (ASA) and its partial charge. In the embedding vector, the 13 first channels were used as a one hot encoding vector for the types of elements that were possibly used according to the SYBYL Mol2 format. An additional channel was used to label $Mg^{2+}$ ions or water molecules.

**GNN architecture**. As many natural things, such as networks, drug-like molecules or larger biomolecules, can be represented as graphs, deep learning applications using GNNs had a significant impact (3). Graphs have an inherent advantage over 3D CNNs, because of their compact representation compared to the sparse representations of 3D CNNs. Moreover, graph-based representations are invariant to rotations and translations, making them a much more natural representation for molecules, as the orientation of the molecule should not affect the data's representation. To represent a neighborhood near $Mg^{2+}$ ion or a water molecule, we represented all the heavy atoms as nodes in a graph. The first node in the graph is always one of these two molecules. Molecules were selected only if they were within 8Å of the RNA atoms. Two nodes are considered connected within the graph if they are at most 8Å apart. In addition to the vector embedding described in the Feature Definition section, each edge also holds the distance between the two atoms. Two main architectures were used in our experiments: graph convolutional networks(4) and graph attention networks (5). Our graph convolutional model consists of multiple graph convolutions followed by batch normalization. The classifier head consists of a global pooling layer named set2set (6) for aggregating the node representations, followed by two linear layers. Our graph attention model uses three attention layers with 12,11,4 heads respectively in each layer. Each graph attention layer is followed by a batch normalization layer. A classification head of two fully connected layers is then used at the end for the classification. A final attempt at creating a model using GNNs was made by combining the two concepts together. First, running the inputs through a few graph attention layers and only then running a convolutional network. As our most successful model used the combination of both types of graph neural networks (Fig. S1).

**Training.** As mentioned in the preprocessing section, an inherent bias was added toward negative samples. This was done purposefully to make our model more sensitive towards positive samples rather than negative samples. To deal with this bias, during training, we used a binary cross entropy loss weighted with a higher weight for positive samples. The loss can be written as such:

$$WeightedBCELoss = -(ylog(p)a + (1 - y)log(1 - p))$$

Where y is the label, p is the prediction, and a is the positive weight multiplied by the positive label loss to combat the positive and negative sample imbalance. We found that a $\sim$ 1.8 worked best for us. All models were trained with either Adam or AdamW optimizers depending on whether or not the model tended to overfit the training data with a learning rate of around 0.0001. Another aspect during training was data augmentation. We tried to add random Gaussian noise to the location of $Mg^{2+}$ ion graphs. This was done to simulate probes in the vicinity of the ion. The data was augmented during training by moving the coordinate of the ion slightly and then computing the distances to the neighboring nodes. The augmentation ensured that no impossible graphs would be created, meaning the $Mg^{2+}$ ion would never be moved too close or too far from one of its neighbors. To our surprise, the model was able to pick up that this noise was being added only to positive samples. We concluded that this noise was interfering with the distribution of some geometric properties of the location of the ion relative to its neighbors.

**Inference.** To generate potential probe positions next to the RNA surface, we calculated the Connolly solvent accessible surface with a probe radius of 1.4Å and density of 0.5 dots per Å. Probe centers are used as potential $Mg^{2+}$ sites. During the inference stage, the model goes over each probe position, creates an embedding of the probe's neighborhood, and predicts whether or not it is a potential $Mg^{2+}$ binding site. To combat the phenomena of many close-together probes near a potential binding site being labeled as positive by the model, we use iterative clustering to select the probe with the highest confidence within a 1.5Å radius.

**IonNet metrics.** In addition to standard metrics for model performance, we also evaluate the accuracy of the predicted positions by our inference pipeline. We define the Distance Center Center metric (DCC) based on the distance between the center of the predicted binding site to the center of the actual binding site as follows:

$$DCC = \frac{\#ions\ with\ positive\ probes\ within \leq 4\text{Å}}{\#of\ ions}$$

**MetalIonRNA comparison**. MetalIonRNA (7) relied on 50 RNA structures, containing 175 $Mg^{2+}$ ions using 5-fold cross-validation. Because this amount of ions is relatively small for training neural networks, we sampled probes (see Inference above) and added them to the dataset. Probes within 0.72Å from experimentally known binding sites of $Mg^{2+}$ were given positive labels, and the rest were given negative labels. We ended up with a training set of 735 positive samples and 39,065 negative samples and a test set with 229 positive samples and 13,571 negative samples in one of our splits. To deal with this data imbalance, we used a random weighted sampler. In our comparison, we reproduced the MetalIonRNA experiment as closely as possible by training our own model on this dataset of RNA structures.

### 2. IonNet performance

**Architectures and model accuracy.** We tested several network architectures, including 3D-CNN, and GNNs with graph attention layers, graph convolution layers, and a combination of both layers (Table S1). 3D-CNN had the worst performance and the longest training time (days on a single GPU) due to suboptimal data representation. GNN-based architectures had a significantly better performance, with the best results achieved by the combination of convolution and attention layers with a training time of a few hours. We believe the reason using attention and convolutional layers together works best is because the attention layers learn intricate connections between nodes, improving the embeddings. Convolutions on the other hand, enforce an inductive bias that, we believe, increases the importance of close-together nodes as multiplication is applied using the distance between neighboring nodes. Four-fold cross-validation was performed with our best-performing model resulting in an even more precise AUROC estimate for the model with a mean of 0.89 (Fig. 2A). We provide the size of each such fold (Table S4, Table S5.)

**ASA is the most important feature for classification.** We performed an ablation study to identify features that contributed most to the model's performance. We removed the charge, ASA, or atom type (Feature definition) for every one of the nodes in the graph by setting their values to zero. We found that the ASA had the most significant contribution to the model's accuracy, followed by the atom type, while the charge had little to no effect (Fig. S8A). We assume that the model focuses on the geometric properties of the structure, more so than the physicochemical ones. ASA values provide information about the shape of the ion neighborhood. Most likely, $Mg^{2+}$ ions are fitted in cavities with sufficient space for them and surrounding waters (Fig. S9). We attribute this to the geometric shape information present in the graph neural network representation. We also assume that the charge information has little contribution because it is redundant due to the atom type.

**Comparison to MetalIonRNA.** MetalIonRNA is a statistical approach for placing ions based on their distances to the N or an O atom pairs. We trained IonNet using the structures from the MetalIonRNA $Mg^{2+}$ dataset. Despite the low number of structures, our model had an average AUROC of 83.3% (Fig. S8.)

**Comparison to MgNet.** MgNet (8) is a 3D convolutional neural network for finding $Mg^{2+}$ binding sites in RNA structures. MgNet's reported metrics over the test set is an average recall of 46.9 and an average precision of 35.5 over all of test set folds, which is significantly lower compared to IonNet. (Fig. 2A, Table S1.)

**Inference scoring with DCC.** Our inference process employs iterative clustering to identify $Mg^{2+}$ positions with the highest confidence, according to IonNet. Subsequently, we evaluated the DCC score on the model's prediction following the iterative clustering step on the $Mg^{2+}$ of our test set. The resulting weighted average DCC score of 0.46 is comparable to the 0.51 recall score exhibited by our model on the test set prior to filtering. This demonstrates that the model's confidence is an effective way to select a subset of $Mg^{2+}$ ion positions without sacrificing much of the model's predictive capabilities (Table S1).
Additionally, we present the distribution of DCC scores across our test set (Fig. S10). Furthermore, we illustrate three examples of our inference outcomes from our test set that are indicative of the model's performance: one exemplary result, one average result, and one poor performing result. These examples were selected from high-resolution structures where the confidence in $Mg^{2+}$ positions is high. These structures are 5nz3 (9) with a resolution of 2.059, 6d8f (10) with a resolution of 2.15 and 6tf2 (11) with a resolution of 2.55. For each structure, we measured a DCC score of 0.6, 1.0, and 0.2, respectively. When increasing the model's confidence threshold for a positive sample from 0.5 to 0.9 the model's DCC on these examples did not decrease (Fig. S11).

**IonNet inference vs Random Selection**:
In Fig. 2 B-D we give an example of IonNet's inference capabilities. Here we expand upon this example and explain the probability of randomly achieving such a result.
In our example (Fig. 2 B-D) there are 5,668 probes, of which only 111 are within a 3Å radius of any $Mg^{2+}$ ion (there are 12 $Mg^{2+}$ ions). The probability of selecting 30 such probes randomly and having none of them within a 3Å radius of any Mg ion is:

$$P(0) = \frac{\left(\frac{5557}{30}\right)}{\left(\frac{5668}{30}\right)} = 54\%$$

Where $\left(\frac{a}{b}\right)$ denotes the combinatorics "choose" operation.
A general* formula for the probability of 'n' correct guesses is

$$P(n) = \frac{\left(\frac{111}{n}\right)*\left(\frac{5557}{30-n}\right)}{\left(\frac{5668}{30}\right)}$$

The expected value is thus.

$$E[correct\ guesses] = \Sigma_{n=0}^{12} P(n) * n \approx 1.58$$

*Note that this is an upper bound on the probability and the expectation since, in this oversimplified calculation, we assume no two probes share the same $Mg^{2+}$ ion as a neighbor

On average, when selecting 30 random probes in this example, we would have only 1.58 good guesses. Whereas in our results, we found 6 such probes, the probability for a random selection such as this is ~0.000018.

**3. Data collection SEC-MALS**

Eluent was subsequently in line with a series of UV at 280nm, MALS, quasi-elastic light scattering (QELS), and refractometer detector. MALS experiments were performed using an 18-angle DAWN HELEOS II light scattering detector connected in tandem with an Optilab refractive index concentration detector (Wyatt Technology). System normalization and calibration were performed with bovine serum albumin using a 55 µL sample at 7 mg/mL in the same SEC running buffer, and a dn/dc value calibrated by BSAS was used to further determine MW by MALS. The light scattering experiments were used to perform analytical scale chromatographic separations for MW determination of the principal peaks in the SEC analysis. UV, MALS, and differential refractive index data were analyzed using Wyatt Astra 7 software to additionally monitor the homogeneity of the sample across the elution peak.

**References:**


1. Berman,H.M. (2000) The Protein Data Bank. *Nucleic Acids Research*, **28**, 235–242.

2. Connolly,M.L. (1983) Analytical molecular surface calculation. *Journal of Applied Crystallography*, **16**, 548–558.

3. Fout,A., Byrd,J., Shariat,B. and Ben-Hur,A. (2017) Protein Interface Prediction using Graph Convolutional Networks. *Adv. Neural Inf. Process. Syst.*, **30**.

4. Rossi,E., Monti,F., Bronstein,M. and Liò,P. (2019) ncRNA Classification with Graph Convolutional Networks. 10.48550/arXiv.1905.06515.

5. Veličković,P., Cucurull,G., Casanova,A., Romero,A., Liò,P. and Bengio,Y. (2017) Graph Attention Networks. 10.48550/arXiv.1710.10903.

6. Vinyals,O., Bengio,S. and Kudlur,M. (2015) Order Matters: Sequence to sequence for sets. 10.48550/arXiv.1511.06391.

7. Philips,A., Milanowska,K., Lach,G., Boniecki,M., Rother,K. and Bujnicki,J.M. (2012) MetalionRNA: computational predictor of metal-binding sites in RNA structures. *Bioinformatics*, **28**, 198–205.

8. Zhou,Y. and Chen,S.-J. (2022) Graph deep learning locates magnesium ions in RNA. *QRB Discov*, **3**.

9. Huang,L., Wang,J., Wilson,T.J. and Lilley,D.M.J. (2017) Structure of the Guanidine III Riboswitch. *Cell Chem Biol*, **24**, 1407–1415.e2.

10. Liu,Y., Esyunina,D., Olovnikov,I., Teplova,M., Kulbachinskiy,A., Aravin,A.A. and Patel,D.J. (2018) Accommodation of Helical Imperfections in Rhodobacter sphaeroides Argonaute Ternary Complexes with Guide RNA and Target DNA. *Cell Rep.*, **24**, 453–462.

11. Huang,L., Wang,J. and Lilley,D.M.J. (2020) Structure and ligand binding of the ADP-binding domain of the NAD riboswitch. *RNA*, **26**, 878–887.

| Model Type | AUROC ↑ | Precision ↑ | Recall ↑ | Specificity ↑ |
|---|---|---|---|---|
| 3D-CNN | 0.73 | 0.68 | 0.40 | 0.90 |
| Graph Attention | 0.86 | 0.79 | 0.50 | 0.82 |
| Graph Convolution | 0.81 | 0.55 | 0.73 | 0.77 |
| Graph Attention + Convolution | 0.88 | 0.82 | 0.51 | 0.96 |

**Table S1.** Network architectures and their performance over all data from the test set, 1371 positive samples and 3584 negative samples

| Sample | SIMPLE SAXS ID | Length (# nucleotide) | Buffer | MW Seq (kDa) | MW SAXS (kDa) | MW MALS (kDa) | Dmax (Å) | Rg (Å) |
|---|---|---|---|---|---|---|---|---|
| #1 | XSEF1RT7 | 28 | 10mM HEPES pH 7.5 100mM KCl, 5mM MgCl2, 1mM TCEP | 8.9 | 10 | ND | 57 | 15.9 |
| #2 | XSSMOQZI | 35 | 10mM HEPES pH 7.5 100mM KCl, 5mM MgCl2, 1mM TCEP | 11.2 | 12 | ND | 60 | 17.4 |
| #3 | XSEFXJZF | 40 | 20mM Tris pH 7.4, 150mM KCl, 5mM MgCl2 | 12.9 | 12 | 12 | 65 | 17.5 |
| #4 | ND | 41 | 20 mM BisTris pH 6.5, 100mM KCl, 2mM MgCl2 | 13.4 | 14 | 19 | 74 | 21.6 |
| #5 | ND | 50 | 20mM Hepes pH 7.4, 100mM KCl, 3mM MgCl2 | 16.0 | 16 | ND | 70 | 21.4 |
| #6 | XSPSCH50 | 66 | 20mM Hepes, 150 mM NaCl, 1 mM TCEP pH 7.5, 2mM Mg Cl2 | 20.5 | 20 | 20 | 108 | 25.8 |
| #7 | ND | 69 | 20 mM BisTris pH 6.5, 100mM KCl, 2mM MgCl2 | 22.3 | 21.4 | ND | 84 | 24.4 |
| #8 | ND | 76 | 20mM Hepes pH 7.4, 100mM KCl, 3mM MgCl2 | 25.2 | 31 | 34 | 100 | 29.6 |
| #9 | ND | 90 | 20mM Hepes pH 7.4, 100mM KCl, 3mM MgCl2 | 29.5 | 33 | ND | 115 | 33.4 |
| #10 | XSIEWNFS | 94 | 20 mM MOPS at pH 6.5, 50 mM KCl, and 7.6 mM $MgCl_2$ + SAM I | 29.1 | 32 | 29.9 | 81 | 22.4 |
| #11 | ND | 111 | 20mM Hepes pH 7.4, 100mM KCl, 3mM MgCl2 | 36.0 | 35.1 | 40 | 160 | 41.1 |
| #12 | ND | 150 | 20mM Hepes pH 7.4, 100mM KCl, 3mM MgCl2 | 48.9 | 42 | 50 | 170 | 41.8 |
| #13 | XSHPX9SN | 160 | 20mM Tris pH 7.4, 300mM KCl, 5mM MgCl2 | 51.8 | 50 | 56 | 108 | 29.6 |
| #14 | XSKSZRJZ | 161 | 20 mM HEPES at pH 6.5, 50 mM KCl, and either 5 mM $MgCl_2$,+2 mM lysine | 49.9 | 47.6 | ND | 108 | 31.1 |

**Table S2.** Experimental SAXS and MALS parameters

| Bin Lengths | Bin Size |
|---|---|
| (0,20) | 130 |
| (20,30) | 112 |
| (30,55) | 110 |
| (55,70) | 97 |
| (70,80) | 92 |
| (80,100) | 102 |
| (100,125) | 149 |
| (125,415) | 91 |
| (450, 2855) | 77 |

**Table S3.** Preprocessing Sequence Bin Sizes

| Dataset | Train Size | Validation Size | Test Size |
|---|---|---|---|
| Fold 1 | 16,482 | 2,445 | 1,123 |
| Fold 2 | 16,601 | 2,334 | 1,062 |
| Fold 3 | 19,855 | 3,324 | 638 |
| Fold 4 | 21,659 | 1,047 | 804 |

**Table S4.** Amount of Mg ion binding sites used in each fold, these numbers do not include the added augmented ion positions later added to increase the size of the dataset.

| Dataset | Positive Samples | Negative Samples |
|---|---|---|
| Fold 1 | 3,177 | 6,405 |
| Fold 2 | 2,869 | 6,977 |
| Fold 3 | 1,371 | 3,584 |
| Fold 4 | 2,147 | 4,342 |

**Table S5.** Amount of positive and negative samples in each test set, this includes the additional augmented data that increases the size of the dataset.

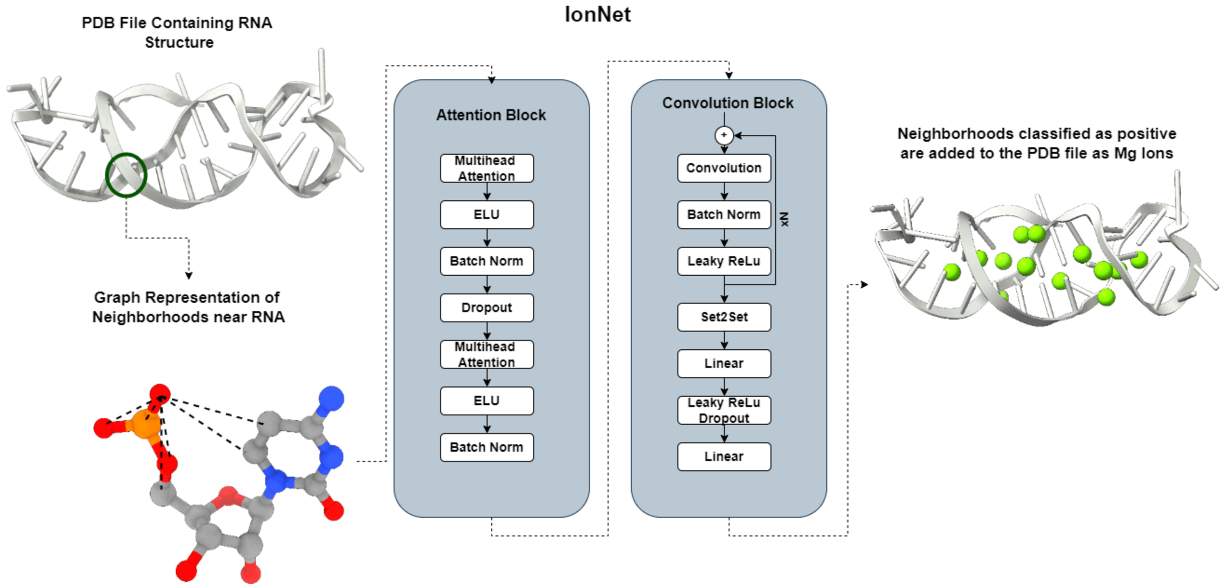


**Figure S1**. Our best performing model architecture, consisting of both graph convolution and graph attention layers.

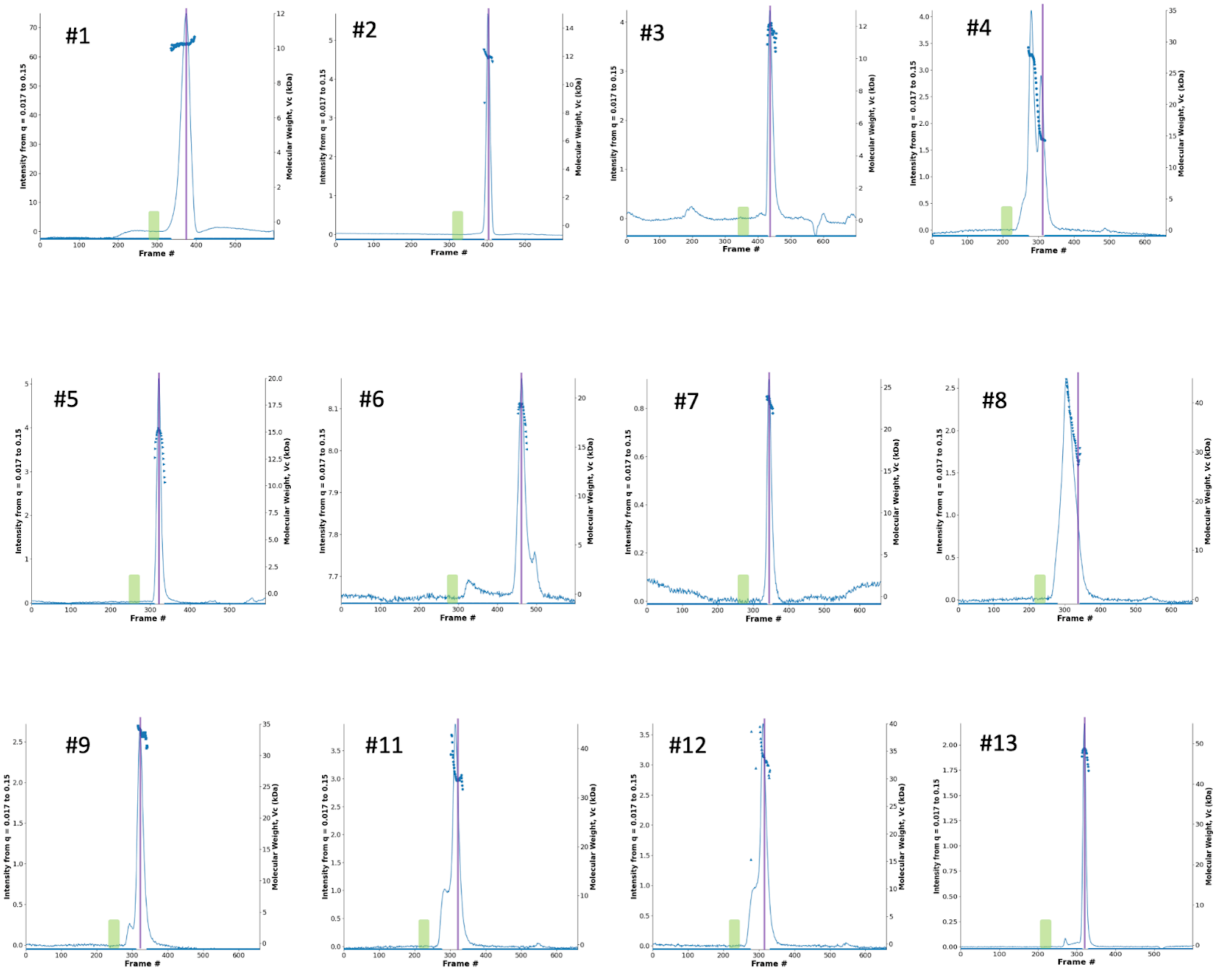


**Figure S2.** The SEC-SAXS chromatograms for each RNA sample show integrated SAXS intensity in the $q$ range of 0.017- 0.17 vs. frame (line), and, if available, Molecular weight vs. frame (symbols). Green-shaded regions are buffer regions, and purple-shaded regions are sample regions. RNA #10 and #14 were collected at the SIBYLS beamline previously and reported in (1,2)

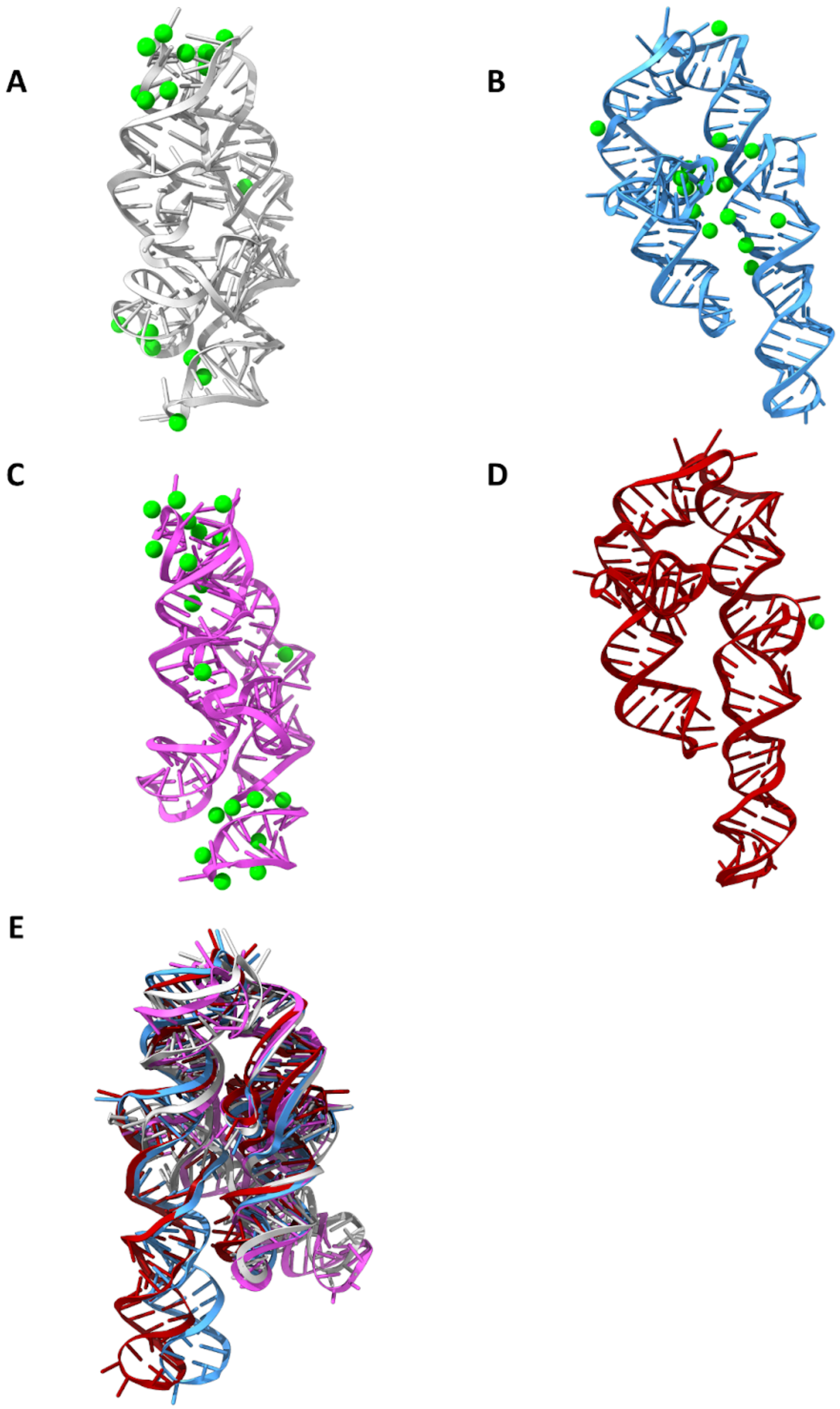


**Figure S3. Multi-state fitting of P4P6 (sample #13)**. A,B,C,D Four conformations were selected, along with their respective ions, by the MultiFoXS program with the weights of 0.084, 0.337, 0.522, and 0.057, respectively. E. Structural alignment of the four conformations using ChimeraX.

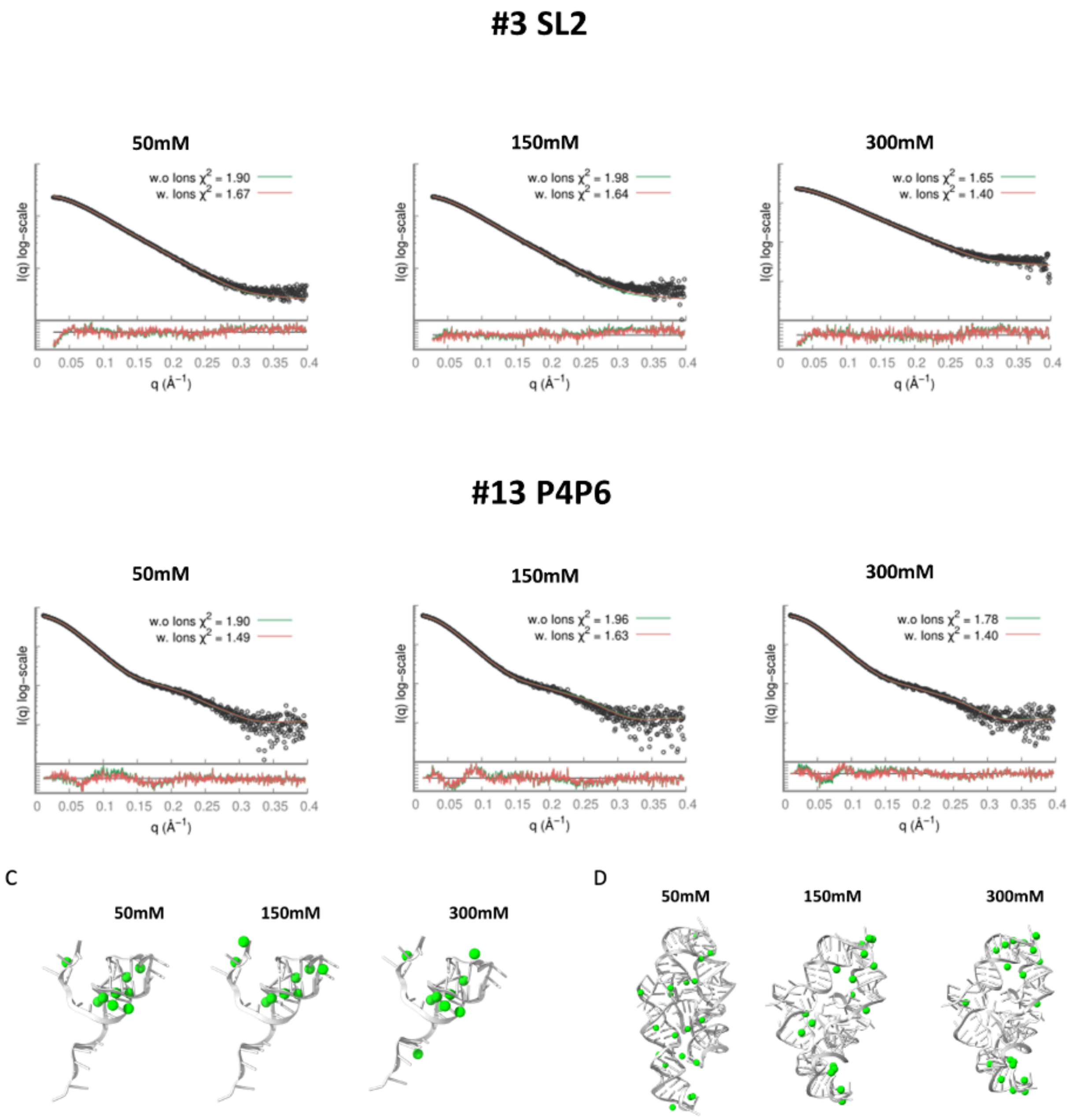


**Figure S4. (A-B)** Experimental SAXS curves (black) for samples #3 and #13 in comparison for the best fit - structure without $Mg^{2+}$ ions and unconstrained c1 and c2 values (green curve). The selected best-fit structure with added $Mg^{2+}$ ions that were selected by running a combinatorial SAXS-based selection of ions while unconstrained. **C-D.** #3 (RNA stem-loop) and #13 (P4P6) structures, respectively. In each figure, we see how ionic strength affects the placement of $Mg^{2+}$ ions (green). The stem-loop structure remains the same across all concentrations, while the selected P4P6's structure at the 50mM KCl shows small differences in the compaction relative to 150mM and 300mM conditions

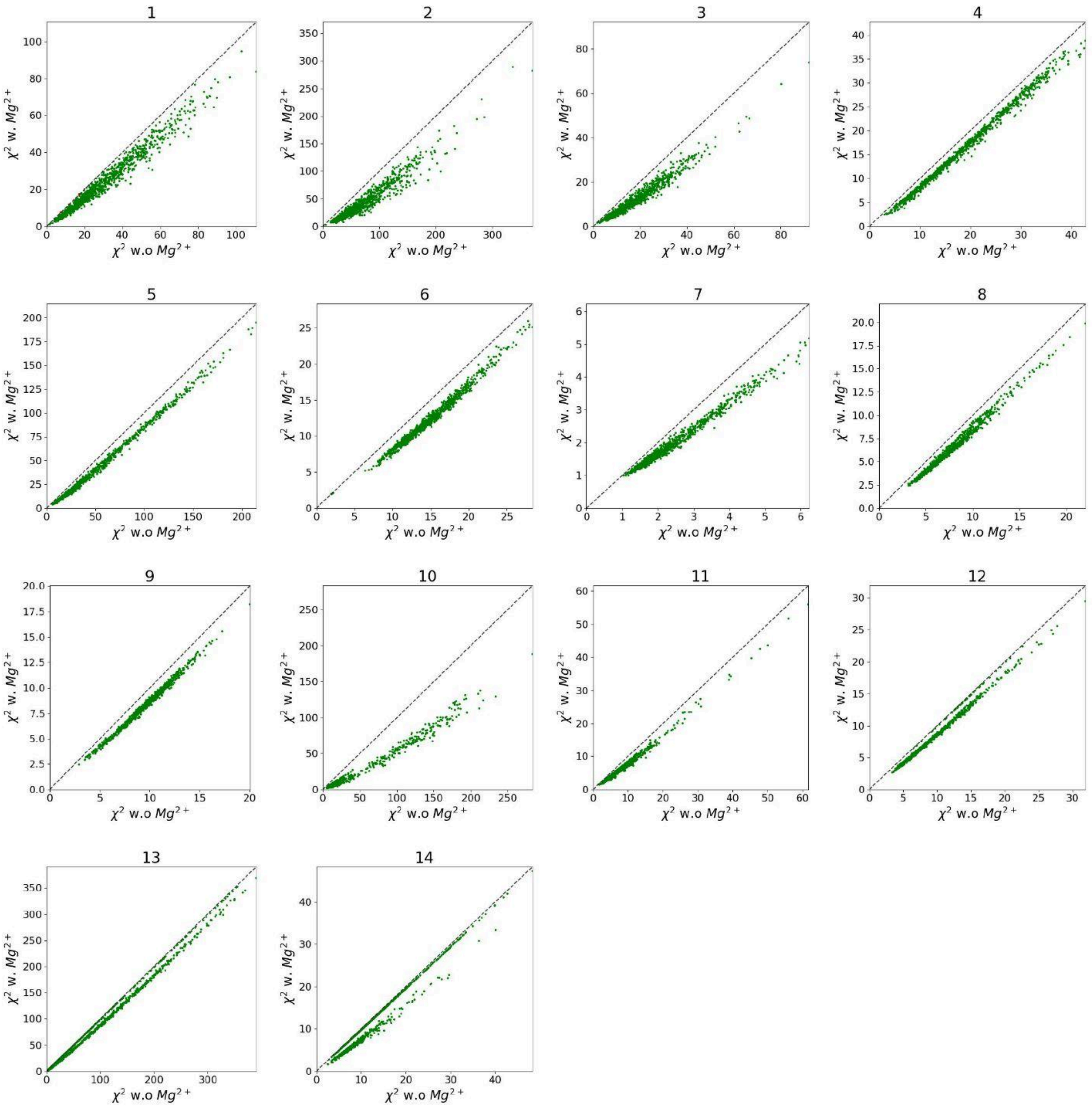


**Figure S5.** $\chi 2$ before and after adding $Mg^{2+}$ ions to all structures sampled by KGSRNA.

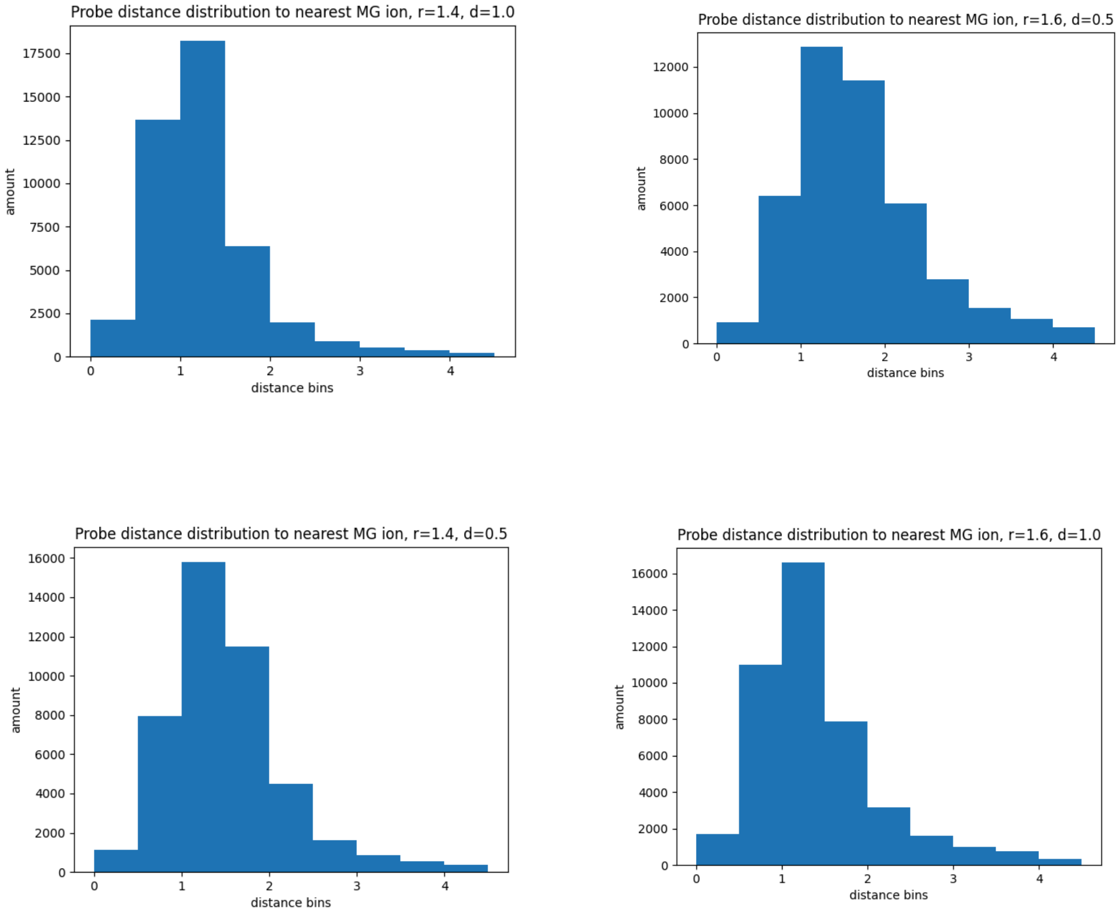


**Figure S6.** Probe distance from ground truth distribution according to different surface probe densities and probe radii of Connolly surface calculation method over all ions in our entire dataset.

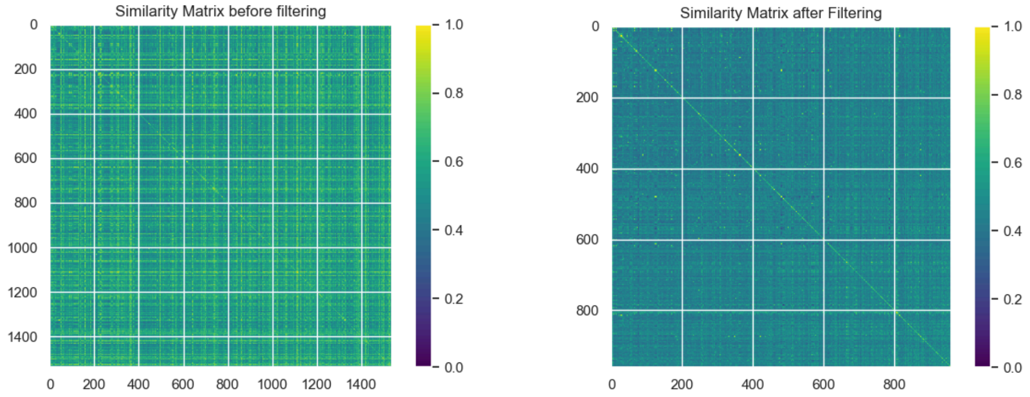


**Figure S7.** Pairwise sequence identity matrix between all RNA chains in the dataset (left) before filtering based on average sequence identity and (right) after filtering.

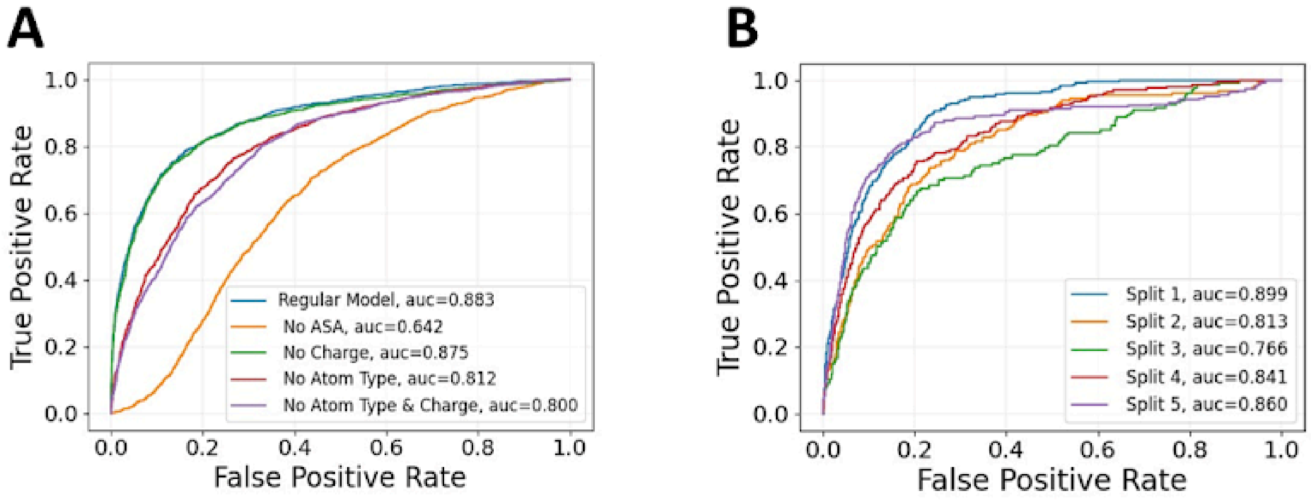


**Figure S8**. IonNet model metrics. A. ROC curves for our best model and ablation studies. B. Results of training using MetalIonRNA dataset.

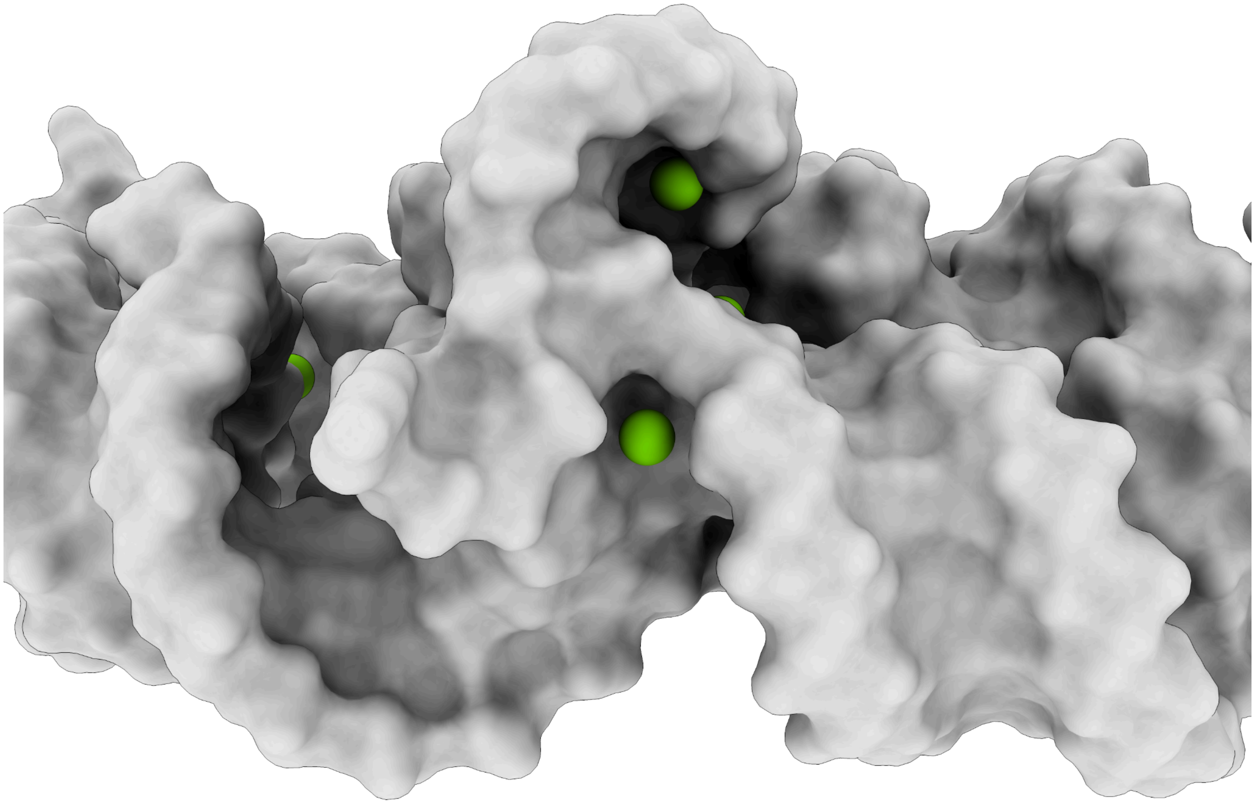

**Figure S9.** IonNet prediction sites in P4P6 structure. We find that IonNet prefers to place $Mg^{2+}$ ions in cavities on the RNA's surface.

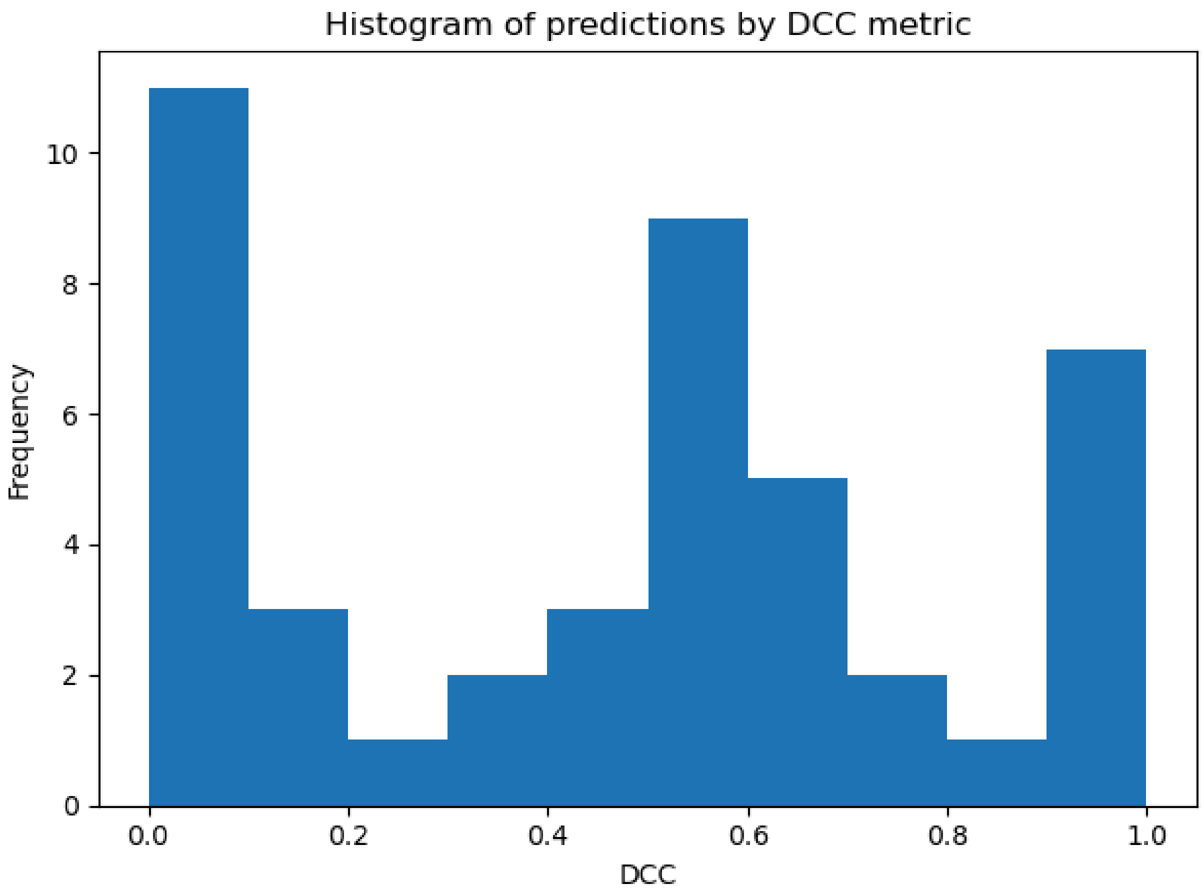


**Figure S10.** Distribution of DCC metric over the test set, with a weighted mean (by the number of $Mg^{2+}$ atoms per test case) of 0.46.

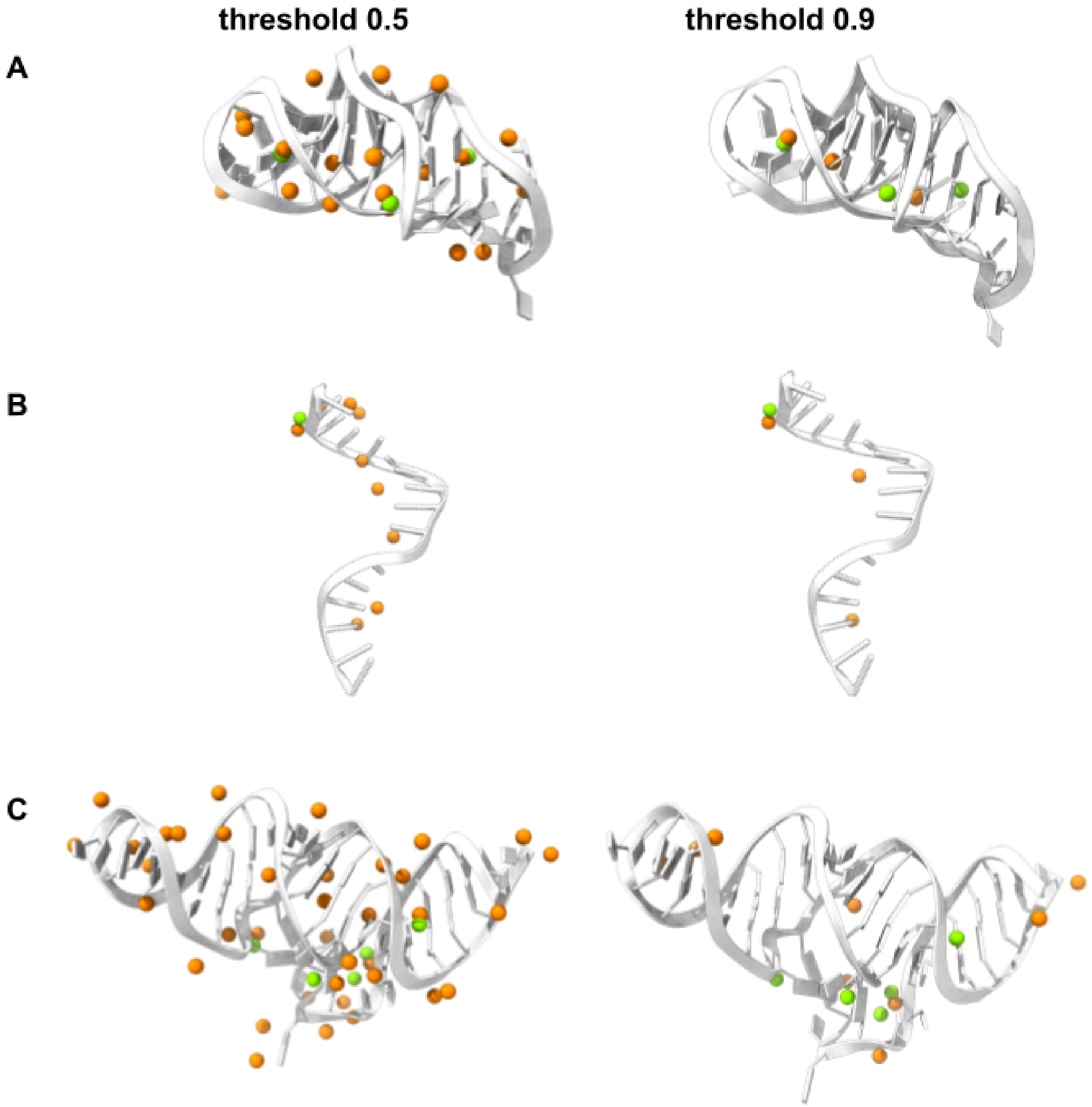


**Figure S11. A.** Guanidine III Riboswitch (PDB 5nz3, resolution 2.059Å), DCC score 0.6. IonNet predicted 18 $Mg^{2+}$ ion positions with a threshold of 0.5 after iterative clustering. Using a threshold of 0.9 resulted in 3 $Mg^{2+}$ ion positions without reducing the DCC score.

**B.** Guide RNA (PDB 6d8f, resolution 2.15Å), DCC score 1.0. IonNet predicted 9 $Mg^{2+}$ ion positions with a threshold of 0.5 after iterative clustering. Using a threshold of 0.9 resulted in 3 $Mg^{2+}$ ion positions without reducing the DCC score.

**C.** Structure of the ADP-binding domain of the NAD+ riboswitch (PDB 6tf2, resolution 2.55Å), DCC score 0.2. IonNet predicted 37 $Mg^{2+}$ ion positions with a threshold of 0.5 after iterative clustering. Using a threshold of 0.9 resulted in 9 $Mg^{2+}$ ion positions without reducing the DCC score.

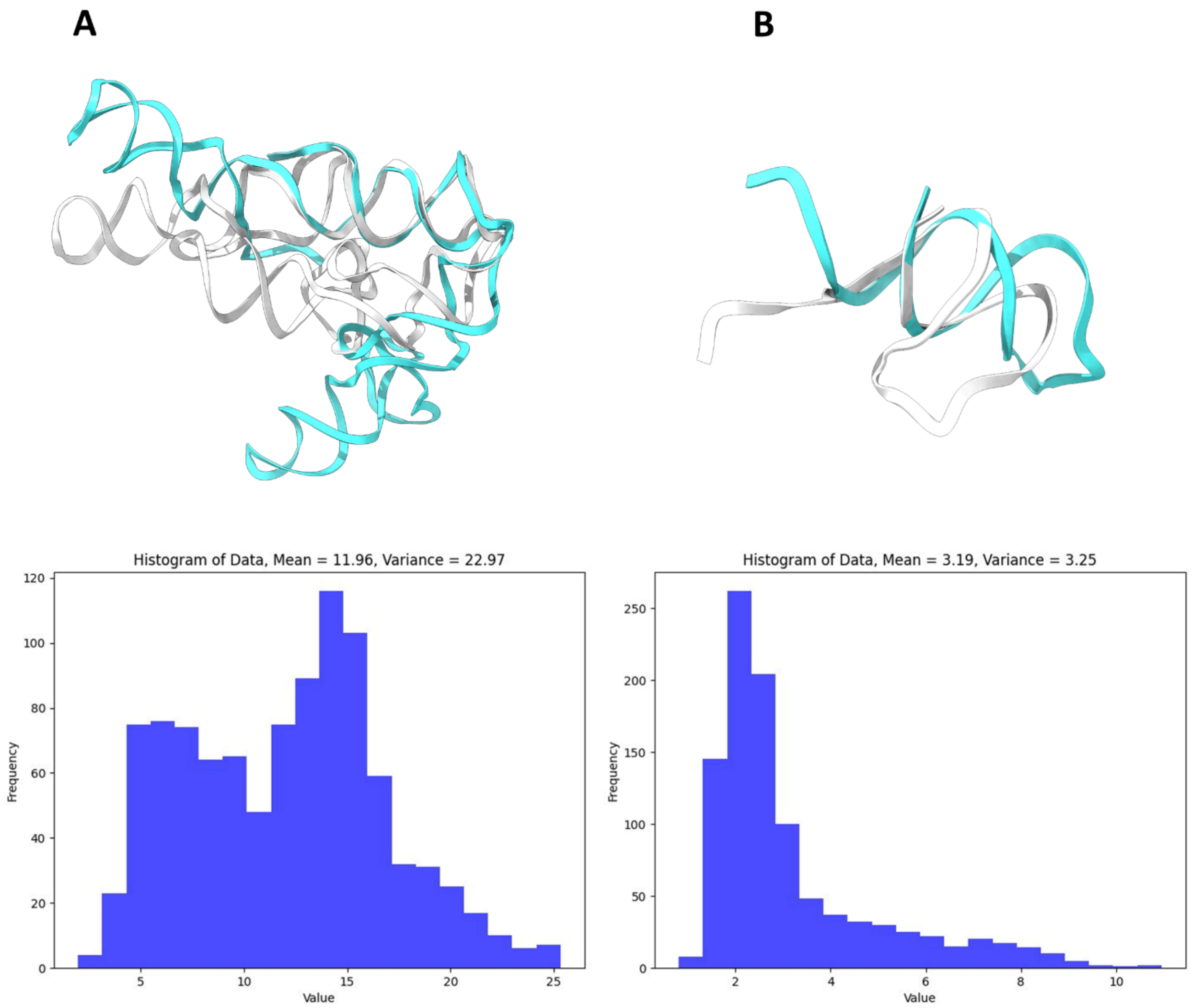


**Figure S12 Column A.** Sample #13 (P4P6) **Column B.** Sample #2 (JK11). Both columns have an image with initial structure (cyan) and the structure sampled by KGSRNA with highest RMSD structure when compared to the initial structure (white). Below the structure images is the distribution of RMSD between the original structure and all of the 1000 samples from KGSRNA.
